# Inward and Outward Tethers Read Out the Spontaneous Curvature of Cellular Membranes

**DOI:** 10.64898/2026.07.31.742037

**Authors:** Beatrice J. Geiger, Luis E. Hamel Ascanio, Effrosyni Drakouli, Victoria Thusgaard Ruhoff, Noëlle Klasner, Younes F. Baroojii, Guillermo Moreno-Pescador, Katrine T. Schjoldager, Hiren J. Joshi, Yoshiki Narimatsu, Signe Mathiasen, Jesper Nylandsted, Poul Martin Bendix, Weria Pezeshkian

## Abstract

Spontaneous curvature characterizes the propensity of a membrane to bend in a specific direction. It is therefore crucial in the multitude of cellular processes that involve membrane shape remodelling. Yet, experimentally quantifying the spontaneous curvature remains a significant challenge in complex biological membranes, as their heterogeneity causes ambiguities in spontaneous curvature’s physical interpretation. Here, we introduce a general experiment-simulation framework to measure an effective spontaneous curvature using dual-direction tether pulling from cell-attached giant plasma membrane vesicles (GPMVs) and mesoscale simulations. For homogeneous membranes, the force difference between inward and outward pulls yields a tension-independent readout of spontaneous curvature. We show that this continuum observable can be generalized to the mean of the spontaneous curvature in a heterogeneous membrane, independent of the underlying microscopic spontaneous curvature distribution. Applied to HEK-derived GPMVs, a baseline negative spontaneous curvature of the plasma membrane is revealed. Sucrose treatment and extracellular addition of Annexin A5 systematically shift the effective spontaneous curvature, while mucin reporter overexpression does not measurably alter it under the conditions tested. We also measure the curvature imprint of individual fluorescently tagged proteins through a sorting index. Benchmarked with Annexin A5, our scheme recovers curvature imprints very similar to previous atomistic molecular dynamics simulations. Taken together this makes spontaneous curvature accessible as a directly measurable material property of native membranes and membrane-proteins, enabling quantitative studies of membrane remodelling across diverse cellular processes.

## INTRODUCTION

Biological membranes are complex thin materials that form the boundary of the cell and compartmentalize eukaryotic cells into specialized organelles. They adopt remarkably diverse shapes, from highly curved endocytic pits and filopodial protrusions to intricate organelle architectures such as endoplasmic reticula, with each morphology supporting distinct cellular functions ^1–3^. One of the main determinants of membrane shape is transmembrane asymmetry, the unequal distribution of biomolecules between the two bilayer leaflets, which has recently become a major focus in cell membrane investigations ^4–7^. At the continuum level, the molecular interactions arising from this asymmetry are collectively captured by the **spontaneous membrane curvature**, *C̅*_0_, a material parameter that quantifies the membrane’s intrinsic tendency to bend. The Helfrich Hamiltonian uses this concept via a bending energy that penalizes local deviations of membrane curvature from *C̅*_0_ ^8,9^. Spontaneous curvature influences how membranes deform, fuse and fissure ^1,10–12^, providing a physical basis for the dynamic architecture of cells and linking effective molecular-scale lipid and protein interactions to large-scale cellular functions.

Despite its central role, spontaneous curvature remains both experimentally challenging to assess and conceptually elusive in complex biological membranes. Only qualitative *C̅*_0_ measurements exist on cell membranes, e.g. shape analyses of red blood cells ^13^ or cell derived vesicles ^14^. More quantitative studies employ model membranes in which spontaneous curvature induced via protein binding or asymmetry of lipids and solvents has been measured ^15–18^. However, a method for quantifying the spontaneous curvature of the cell’s plasma membrane expressing various endogenous curvature active proteins is urgently needed for precise prediction of membrane remodelling events and the sorting and function of curvature-inducing proteins.

A common method for measuring macroscopic mechanical properties of a material is to perturb it from a relaxed state and quantify its response. For membrane systems, tether pulling provides an effective scheme: for an idealized geometry pulled from a homogeneous membrane, the tether’s mechanics depend on the membrane bending rigidity *κ*, surface tension *τ*, and spontaneous curvature *C̅*_0_. The spontaneous curvature of synthetic Giant Unilamellar Vesicles (GUVs), with controlled tension and well-defined bending rigidity, can be quantified by employing optical tweezers to extract membrane nanotubes (pull outward) while quantifying the tether’s holding force. As an interesting development, tubes have also been pulled into GUVs ^18^ thus generating membrane curvature of opposite sign, similar to membrane invaginations present in endocytic buds or cristae in mitochondria. An especially important theoretical relation arises when two tubes with opposite curvature are pulled consecutively from the same membrane. Assuming a homogeneous membrane, the difference in the holding force is then directly proportional to the spontaneous curvature (see Ref. ^18^ and Supplementary Note 1),

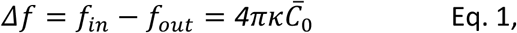

where *f_in_*, *f_out_* are the holding forces for inward- and outward-pulled tethers, respectively. This allows quantification of spontaneous curvature independent of the membrane tension as long as the tension stays constant during both pulls. Despite the potential of such dual-tether pulling, this approach has so far been unexploited for exploring the effects of integral membrane proteins and of peripheral binding proteins on the spontaneous curvature of membranes. Experimentally, quantifying spontaneous membrane curvature in cellular membranes rather than in synthetic vesicles like in Ref. ^18^, provides several benefits: i) membrane proteins can be expressed with fluorescent tags in the donor cell; ii) transmembrane proteins keep their natural orientation (in→out) in the membrane; and iii) curvature effects can be explored in the native environment directly relating a physical parameter to a biological system.

However, in applying the idealized theory to cellular membranes, one encounters two main confounders: the influence of membrane fluctuations, especially at low membrane tensions, and the presence of multiple curvature-inducing protein- and possibly lipid-types meaning spontaneous curvature is no longer spatially uniform (we refer to this as a heterogeneous membrane). While the first issue can be addressed by fluctuation corrections appended to simpler formulae ^19^, the second problem persists, demanding at least one additional degree of freedom to be added to the Helfrich Hamiltonian.

In the present work we propose a solution to measure the spontaneous curvature of cellular membranes and the spontaneous curvature imprint of individual proteins by employing dual-tether pulling experiments on cell-attached giant plasma membrane vesicles (GPMVs) combined with mesoscale simulations ^20^ to interpret the experimental results. We established a tether pulling protocol suitable to measure the force difference between out- and in-pulls on the cell detached plasma membrane of mammalian cells (Figure 1). To understand the physics governing the deformation of heterogeneous membranes we perform tether pulling in mesoscale simulations. After validating that our simulation scheme captures the results of standard tether pulling theory on homogeneous membranes, we extend the Helfrich Hamiltonian to a heterogeneous membrane by introducing a scalar local spontaneous curvature field. In this framework, each simulated membrane segment is assigned a distinct preferred curvature value *c*_0_ drawn from a distribution with mean ⟨*c_0_*⟩ and the field is free to reorganize on the membrane surface. Based on the mesoscale simulations, we construct a link between the force difference in dual-tether pulling and the mean of the heterogeneous spontaneous curvature. This provides a systematic tool for mapping experimental observations onto the theoretical landscape, enabling the extraction of mean spontaneous curvature for different complex membrane systems. We propose curvature-induced sorting of inclusions as an additional measure that can even provide insights to individual protein species’ curvature imprint on native membranes.

**Figure 1:**
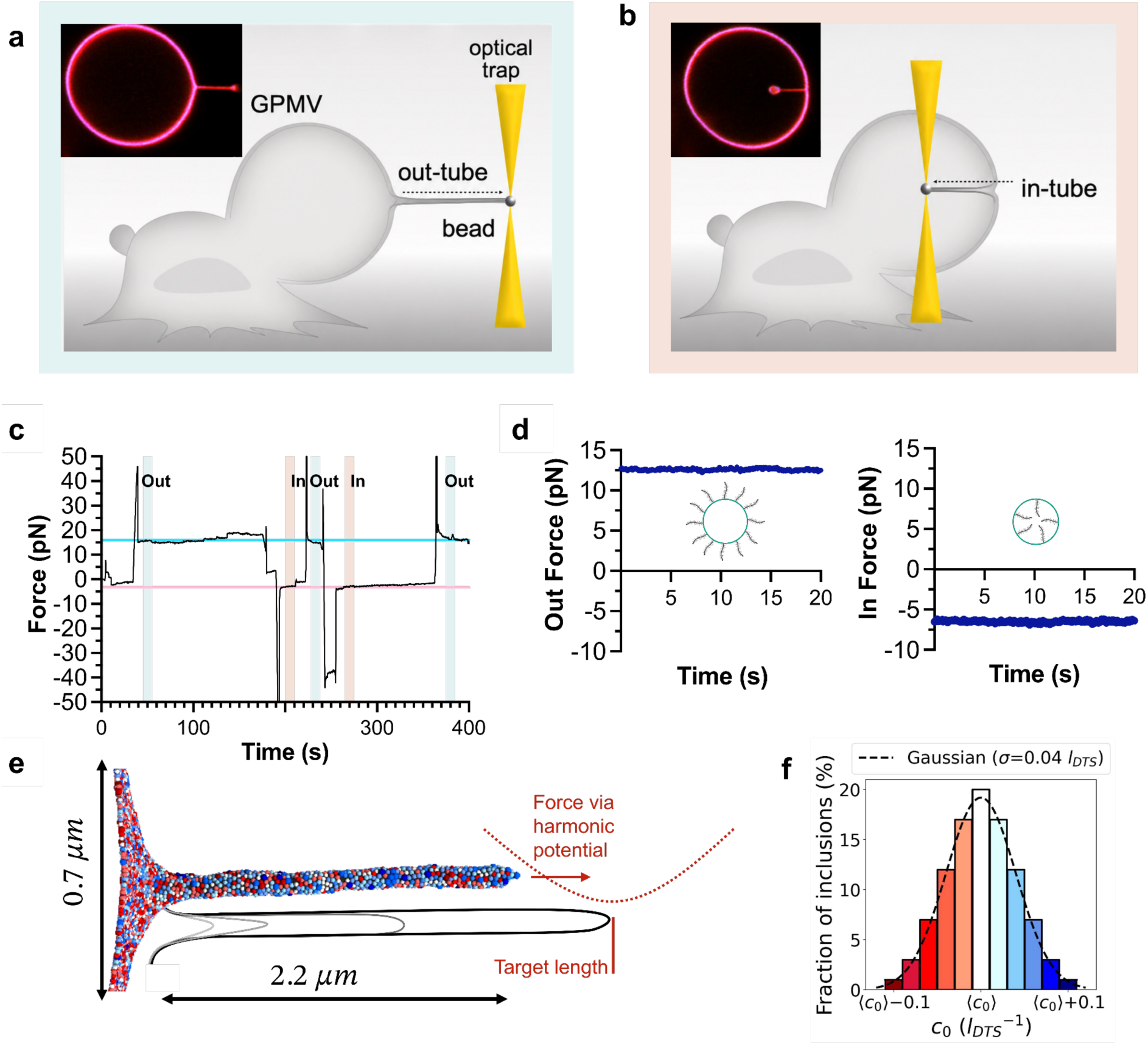
Dual membrane tether pulling from GPMVs using optical tweezers and with FreeDTS simulations. (a&b) Schematic of an out-tube (a) or in-tube (b) configuration where a bead attached to the GPMV is pulled via an optical trap, forming a membrane nanotube into the extracellular space or as an invagination respectively. Insets show the corresponding confocal microscopy images of a GPMV attached to its parent HEK293 cell, with an overlay of membrane label DiD (red) and expressed GFP-Mucin-5AC (blue). **(c)** Force vs. time plot for a successive dual-tether measurement. Turquoise and red shading denote the start of a plateau after the formation of out-tubes and in-tubes, respectively. Horizontal lines indicate average force plateau values. Transient force peaks observed at the start of each pull correspond to the active deformation phase prior to nanotube formation and are excluded from analysis. The consistency of force plateaus across multiple pulls confirms constant vesicle tension. **(d)** Representative force vs. time plot for the out-pull plateau (left) or the in-pull plateau (right) used for analysis. Inset: schematic of a cross-section of tethers, where mucin N-termini extend into the extracellular space. **(e)** Simulation setup for simulated tether pulling of a membrane decorated by 100% of inclusions which locally induce different spontaneous curvatures *c_0_* represented by different colors (see (f)), yielding a mobile heterogeneous spontaneous curvature field. The tether is pulled to a target length via a harmonic potential between a chosen tip vertex position and the remaining vertices’ positions. Conversion to physical units here was performed with respect to HEK-cell controls (see Methods). **(f)** Visualisation of an exemplary gaussian inclusion distribution with 11 inclusion species, representing for example proteins of different preferred spontaneous curvature.

## RESULTS

### 1.1 Force measurements on dual tethers of cell membranes

Cultured cells were firmly adhered to a poly-L-lysine (PLL) coated substrate and NEM (N-ethyl maleimide) was added to trigger the cell membrane to detach from the underlying actin cortex to form cell-attached GPMVs (Figure 1a&b). NEM is known not to cause cross linking of membrane proteins unlike other methods using DTT ^21^. This procedure allows us to produce cell attached vesicles that retain the plasma membrane complexity to a large extent. Notably, protein orientation is preserved ^22^ and the cell-attached GPMVs are devoid of larger intracellular components, such as cytoskeletal elements, which allows the isolated study of membrane-associated processes in the absence of active cellular remodelling dynamics.

Membrane nanotubes were extracted from these cell-attached GPMVs both outwards (Figure 1a) and inwards (Figure 1b) while the force and fluorescence intensity were recorded. Inward pulling requires careful focal adjustment to keep the nanotube in focus (Supplementary Figure S1). The dual tube pulling causes the membrane to form opposite curvatures, with the external membrane leaflet of the cell on the tube’s outer surface during the out-pull, see Figure 1d (left panel), and on the tube’s inner side during the in-pull, see Figure 1d (right panel). The corresponding outward force magnitude, *f* = 13*pN* is then subtracted from the required inward force magnitude *f* = 5*pN* (Figure 1d) resulting in Δ*f* = −8*pN*. The use of in- and outward forces to obtain the spontaneous curvature of the membrane requires constant membrane tension during both pulls, which we verify by performing multiple subsequent pulls on the same GPMV in both directions (Figure 1c). We consistently obtain similar forces.

The forces measured as shown in Figure 1d then suggest a spontaneous curvature −0.012 *nm*^−^*^1^*≤ *C̅*_0_ ≤ −0.004 *nm*^−^*^1^* (Supplementary Figure S2) using Equation 1 and published values for the bending stiffness of plasma membranes (10 − 30*k_B_T* see Ref. ^23–26^) in the idealized homogeneous membrane model.

### 1.2 Dual-tether simulations highlight advantages of force difference - spontaneous curvature relation

To correctly exploit the dual-tether pulling assay on GPMVs one must replace the laterally constant parameter *C̅*_0_in Eq. 1 with a quantity which represents their heterogeneous spontaneous curvature. For this we introduce in-silico mesoscale tether pulling using the FreeDTS software ^20^ that models a membrane as a dynamically triangulated surface. Membrane bending energy is governed by a discretized Helfrich-Hamiltonian and equilibrium configurations are sampled using a Monte Carlo algorithm (see Methods). The simulation model uses an internal length scale, *l_DTS_*, which can be converted to physical units for the biological system of interest (see below, Methods, and other examples in Ref. ^20^).

We simulate tether pulling experiments on a flat membrane patch in periodic boundary conditions (PBC) (Figure 1e, Methods), under constant externally applied frame tension *τ* ^27^. This ensemble is shown to reproduce membrane tension under theoretically tractable conditions ^28^ (see Methods). We directly control the membrane bending rigidity *κ*, and the target tether length *L*. Compared to higher resolution models, e.g. molecular dynamics simulations, the mesoscale approach enables the desired time and length scales to determine the tether’s equilibrium properties.

As a prerequisite one must find those target tether lengths *L* at which standard tether pulling theory becomes applicable to the simulations. For this we monitored the ratio of a measured extrusion force to the theoretically expected extrusion force as a function of *L* on a homogeneous membrane (*C̅*_0_ = 0, *τ* = 1.5 *k_B_T*/ *l_DTS_*^2^, *κ* = 20 *k_B_ T*). Such a profile reveals when the saturation regime is reached where radius and holding force fulfil (derivation Supplementary Note 1)

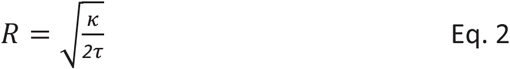

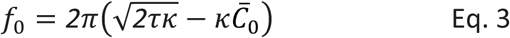

with *C̅*_0_ = 0. This describes the experimental regime in which a membrane reservoir facilitates the elongation allowing the force to be constant over a wide range of tether lengths ^29^ (see also Figure 1c). Our force reached a plateau for tethers corresponding to *L* > 35*l_DTS_* at 70% - 80% of the theoretically expected force (Supplementary Figure S3), in agreement with reports from other mesoscale simulations ^30^. We applied the fluctuation correction 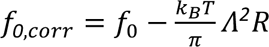 (Ref. ^19^) to the theoretical force, fitting the cutoff wavelength *Λ* to our plateau region. The recovered value for *Λ* agrees with the lower resolution limit of FreeDTS, confirming that forces on simulated tethers closely follow the theory for homogeneous membranes (Supplementary Figure S3 b).

On tethers with *L* = 60*l_DTS_* safely within the force plateau, we next tested the effect of uniform, homogeneous spontaneous curvature in our model (Eq. 3 with *C̅*_0_ ≠ 0). We found good qualitative agreement between measured simulation forces and theory (Supplementary Figure S4), but quantitatively, *Δf* agreed better with theory than *f* under non-zero spontaneous curvature by partially compensating the effect of membrane fluctuations (for details see Supplementary Note 2, Supplementary Figure S5).

To compare simulated to experimental systems going forward, in principle, membrane tension can be used to convert *l_DTS_* to a physical unit. Since tension may vary between different experiments and we do not directly control it (only required to be constant within individual experiments), this conversion would be specific to each. Instead, we use a convenient rescaling of force differences and curvatures by the sum of the in-pulling and out-pulling force (see Methods), enabling direct comparison of all simulation and experimental results. The simulated rescaled force difference *Δf*/(*f_in_* + *f_out_*) as a function of *C̅*_0_/(*f_in_* + *f_out_*) shows the same level of agreement with theory as the scaling-free force difference described above (Supplementary Fig. S4b).

### 2.1 Spontaneous curvature of the cell membrane

Although the plasma membrane is a highly complex system in which spontaneous curvature is unlikely to be uniform, the force difference between dual tether pulls can be related to an effective parameter that captures its net behaviour as *Δf* = *4πκC_0eff_*. We will validate this and provide a meaning of *C_0eff_*, in terms of the local spontaneous curvature distributions representing membrane compositions, in section 2.3. We now apply the experimental approach described in Section 1.1, using the force difference on HEK293 cells to extract the effective spontaneous curvature (*C_0eff_*) of their plasma membrane.

We consistently find that the force magnitudes required to pull tubes outwards (Figure 2a) are higher than for extraction of inward tubes (Figure 2b). This suggests a negative spontaneous curvature of wild-type HEK293 cell membranes labelled with a DiD membrane marker. Their effective spontaneous curvature at *κ* = 20*k_B_T* (see Ref. ^21^) lies between −12.9 × 10^−3^*nm*^−^*^1^* and −0.48 × 10^−3^*nm*^−^*^1^*(Figure 2c).

**Figure 2:**
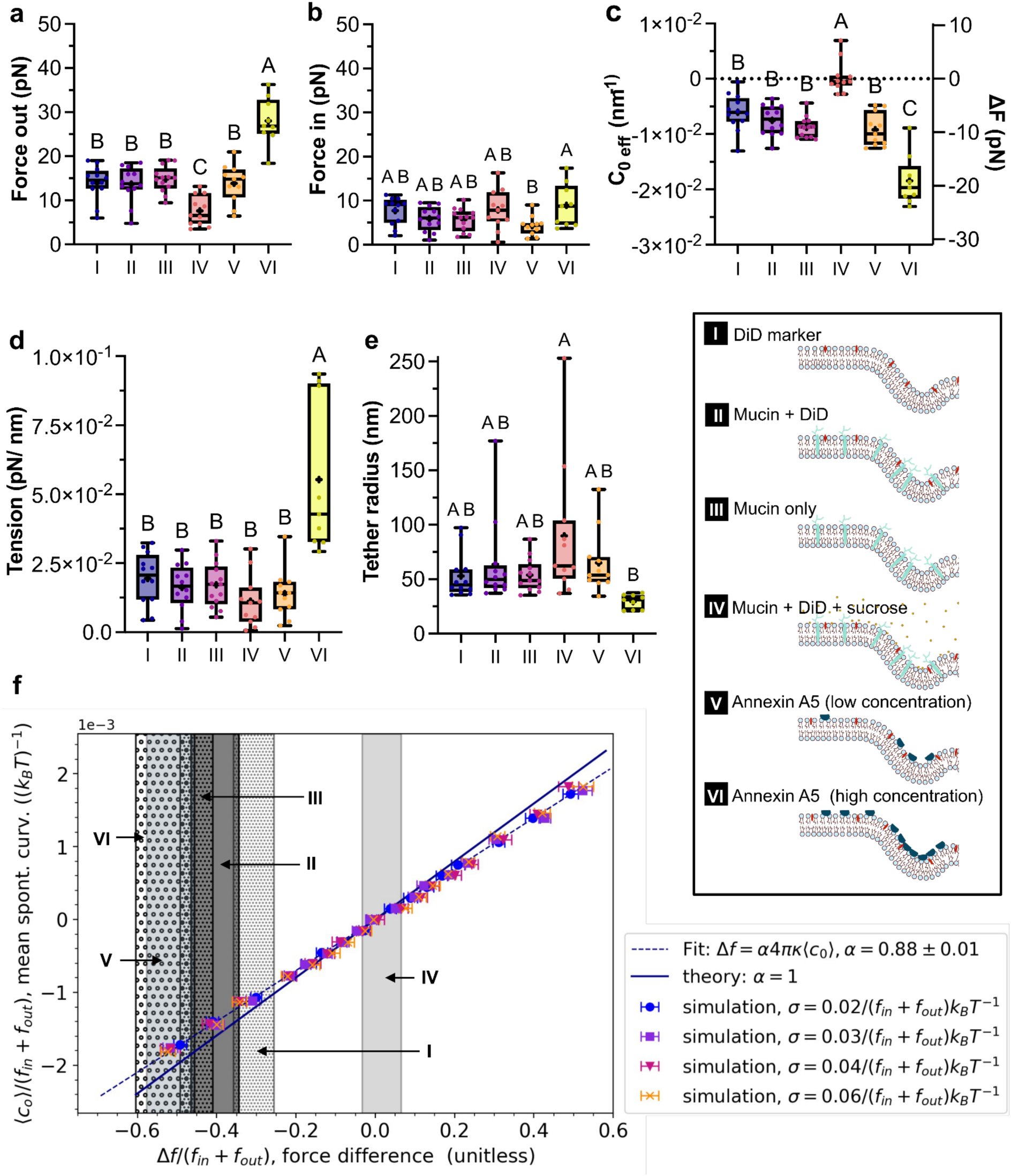
Membrane mechanics and spontaneous curvature measured via dual tether pulling on GPMVs and simulated membranes with heterogeneous inclusion mixtures. (a&b) Box plots showing the average force required to pull membrane tubes either outward (a) or inward (b) from GPMVs derived from HEK293 cells under different conditions. **(c)** Box plots showing the effective spontaneous curvature, calculated using equation 1 in the form *Δf* = 4*πκC_0eff_*. **(d)** Membrane tension calculated from force measurements using equation S1-6. **(e)** Tether radius calculated using equation 2, with the tension values shown in (d). The bending modulus was assumed to be *κ* = 20*k_B_T* = 82.3 *pNnm* in (c), (d), (e). Each box represents the interquartile range (IQR), with the median and mean (+) indicated. Groups with different capital letters differ significantly (P<0.05), whereas groups sharing the same letter do not (ordinary one-way ANOVA with Tukey’s multiple-comparisons test). Number of experimental days, n, and datapoints, N, in (a)-(e): n_I_ = 4, N_I_ = 12; n_II_ = 5, N_II_ = 12; n_III_ = 5, N_III_ = 13; n_IV_ = 2, N_IV_ = 11; n_V_ = 5, N_V_ = 11; n_VI_ = 3, N_VI_ = 9. **(f)** Difference in pulling force in simulated tethers at *κ* = 20*k_B_T* with a heterogeneous curvature field represented by an inclusion mixture of 11 different inclusion species. Membranes were fully covered (1 inclusion per vertex). The inclusion species differ by their curvature preference with respect to the mean ⟨*c_0_*⟩ of the mixture (see also Fig. 1f). The abundance of each species is chosen so that the population approximates a normal distribution with mean ⟨*c_0_*⟩ and standard deviation *σ*. Each datapoint represents the analysis of the last 2 × 10^7^ MC steps from 5 replica of equilibrated simulations (run for 12 × 10^7^ MC steps), error bars indicate the SEM obtained via block averaging.

We examined cell membranes expressing fluorescent mucins as potentially curvature-inducing integral transmembrane proteins. Mucins are characterized by O-glycosylated tandem repeat (TR) domains and overexpression of mucin 1 (MUC1) can induce outward membrane bending through steric interactions amongst the glycopolymers ^31^. To examine the effect on membrane mechanics, we used a highly glycosylated mucin 5AC TR (GFP-MUC5AC) with a MUC1-derived transmembrane anchor (see Methods) ^32^. The results show no increase in effective spontaneous curvature by MUC5AC at our expression levels, compared with wild-type HEK293 cell controls (Figure 2a-c). Our truncated MUC5AC construct was shorter than the full-length MUC1 used in Ref. ^31^ which may have reduced steric crowding. Additionally, heterogeneous MUC5AC expression levels among different cells (Supplementary Figure S7) complicated population-level measurements of any potential crowding effect. We observed several select GPMVs displaying internal buds (Supplementary Figure S6), supporting that MUC5AC cell membranes retain a negative spontaneous curvature.

To examine changes in spontaneous curvature induced by external perturbations of the membrane, we subsequently replaced the extracellular medium with a solution containing sucrose. High concentrations of sugars are known to expand the proximal leaflet resulting in increased spontaneous curvatures ^33^. Adding 500 mM sucrose to cells expressing MUC5AC accordingly caused a statistically significant increase of the effective spontaneous curvature at *κ* = 20*k_B_T* from (−7.6 ± 0.8) × 10^−3^*nm*^−^*^1^*without, to (−0.1990 ± 0.7) × 10^−3^*nm*^−^*^1^* with sucrose (Figure 2c).

To test if our method could detect spontaneous curvature decreases from binding of a curvature modulating protein, we extracellularly added recombinant Annexin A5-AF488 that binds to the external leaflet containing phosphatidylserine lipids in blebbing cells. Addition of Annexin A5 lowered the effective spontaneous curvature in a concentration-dependent manner. Through addition of 5*μL* Annexin A5 probe (see Methods), denoted as low concentration, a decrease to *C_0eff_* = (−9.2 ± 0.9) × 10^−3^*nm*^−^*^1^* was found, while addition of 50*μL* Annexin A5 probe, denoted as high concentration, resulted in a statistically significant decrease to (−18.5 ± 1.5) × 10^−3^*nm*^−^*^1^*(Fig. 2a-c). The dependence on actual membrane-bound protein concentration was recovered using protein-channel associated fluorescence at the GPMV surface, confirming a surface density-dependent *C_0eff_* modulation (Supplementary Figure S8).

### 2.2 Cell attached GPMVs have low membrane tension

From the force sum *f_in_* + *f_out_* acquired in our two-way tether assay we determined the membrane tension (Supplementary Note 1, Eq. S1-6). As shown in Figure 2d the tensions in the cell attached GPMVs lie in the range ≈ 0.005 − 0.02 *pN*/*nm*, except for high Annexin A5 concentrations. This is a low-to-medium tension regime, compared to the higher tensions of ≥ 0.016 − 0.064 *pN*/*nm* used in previous dual tether pulling assays on synthetic model membranes ^18^. We therefore not only capture native plasma membrane compositions but also mechanical environments closer to membrane tensions in living cells ^34–36^ and far lower than the membrane rupture tension of ≈ 1 − 10 *pN*/*nm* ^37–40^.

The radii of the nanotubes could be calculated based on the tension recovered from force measurements and mostly lay in the range of 50 − 150 *nm* (Figure 2e). According to Eq. 2 both inward and outward tubes pulled from the same membrane, having constant membrane tension *τ*, will have the same radius even at non-zero spontaneous curvature (derivation in Supplementary Note 1). This was validated through additional measurements of the radii via the fluorescent membrane marker signals (linearly proportional to tube radius) and comparing tether sizes through pairwise recordings of in- and outward tethers as shown in Supplementary Figures S9 and S10. We note that the latter approach is however sensitive to bleaching and out-of-focus imaging artefacts between the two tether images.

### 2.3. In complex membranes ***Δf*** robustly determines the mean of heterogeneous spontaneous curvature

To interpret our experimental results, we needed to connect the extracted effective curvature values *C_0eff_* to the underlying membrane composition. For this we returned to the mesoscale simulations and probed the relation of *Δf* to the spontaneous curvature distribution of heterogeneous membranes. Again, the in- and out-tether pulling experiment was modelled on a flat membrane patch under PBC (bending rigidity *κ* = 20*k_B_ T*, tension *τ* = 2 *k_B_T*/*l^2^_DTS_*, target tether length *L* = 60 *l_DTS_*). The model regulates membrane complexity by extending the Helfrich-Hamiltonian with a scalar, local spontaneous curvature field. Each vertex of the simulated membrane, i.e. each membrane segment, is covered with an inclusion. An inclusion of species *i* imposes a local spontaneous curvature *c_0_*_,*i*_ and can represent a curvature preferring protein, asymmetric lipid patch or other molecular agents causing local membrane curvature. We composed the heterogeneous membrane of eleven inclusion species *i* = 0,1,2, …, 10. The abundance of each species approximates either a gaussian, uniform or asymmetric distribution with mean ⟨*c_0_*⟩, and the preferred curvature imposed locally by an individual of the species *i* is *c_0_*_,*i*_ = 0.02 × *i* + ⟨*c_0_*⟩ − 0.1 *l_DTS_*^−^*^1^* (see Fig. 1f). For example, in the distribution around ⟨*c_0_*⟩ = 0 the local curvatures [−0.1, −0.08, −0.06, …, 0.06, 0.08, 0.1] *l_DTS_*^−^*^1^*converted for the control experiments on HEK cells (see Methods) correspond to [−4.9, −3.9, −2.9, …, 2.9, 3.9, 4.9] × 10^−3^*nm*^−^*^1^*.

We performed simulations with these heterogeneous mixtures for varying mean ⟨*c_0_*⟩ and variance *σ* of the distributions. The individual inclusions were randomly distributed on the tethered membrane at the start of each simulation, after which the system was equilibrated, letting both membrane shape and curvature field positions evolve.

We then examined the difference in rescaled pulling force *Δf*/(*f_in_* + *f_out_*) as a function of rescaled mean spontaneous curvature ⟨*c_0_*⟩/(*f_in_* + *f_out_*) of the inclusion distribution. Across a wide range of ⟨*c_0_*⟩ the effective curvature *C_0eff_* that would be extracted like in Eq. 1, closely follows the mean of the curvature field for all gaussian distributions with different variances *σ* (Fig. 2f), as well as uniform and asymmetric distributions (Supplementary Figure S11a,b). Furthermore, we cover the regions where our experimental measurements (section 2.1) lie, see Fig. 2f, showing that *Δf* = 4*πκ*⟨*c_0_*⟩ holds for these membrane compositions in particular for small absolute ⟨*c_0_*⟩ With increasing absolute spontaneous curvatures an effective *C_0eff_* extracted from *Δf* through Eq. 1 diverges from ⟨*c_0_*⟩. However, the mesoscale simulations provide a reliable link allowing for simple corrections to either the slope in Eq. 1 or direct readoff of ⟨*c_0_*⟩ from a plot like Fig. 2f after experimental force measurement. One should note that the data presented here applies to bending rigidities within the biologically relevant regime, but for *κ* >> 20*k_B_ T* or *κ* << 20*k_B_ T* the relation between forces and bending rigidity may change due to *κ*’s effect on fluctuations.

### 3.1 Curvature-induced sorting on GPMV dual tethers

Membrane curvature can function as a biochemical cue in living cells by directing the localization of remodelling machinery, which has been investigated in vitro for example through measuring curvature sorting parameters ^41–45^. For peripheral proteins, the binding affinity to the membrane and further localization depends on the curvature of the leaflet the protein faces. Notably, some unconventionally secreted proteins drive different functions when facing the cytosolic or the exoplasmic leaflet. For example, cytosolic Annexin A5 promotes membrane repair, while extracellular Annexin A5 shields procoagulant membranes ^46,47^. Previous efforts to measure curvature sensing in both curvature directions focused on minimal model systems like patterned platforms or tether-pulling on GUVs ^48,49^. In comparison, measuring the same protein’s sorting to both positive and negative curvature under identical native plasma membrane composition and matched surface density would be advantageous, because (i) many proteins both sense and generate curvature in a concentration-dependent way ^50^, and (ii) comparisons across opposite-curvature assays are easily confounded by different leaflet access, adsorption kinetics, etc.

Hence, we extended our dual tether pulling assay to measure the sorting of different components and used the mesoscale model to connect sorting to individual protein curvature imprints. In the experiment, we compared the signal intensity of fluorescently tagged proteins of interest on tethers, *I_prot_*_−*tet*ℎ*er*_, to the approximately flat GPMV membrane, *I_prot_*_−*flat*_. From these we extracted an experimental sorting index ^51^ on in- or out-pulls:

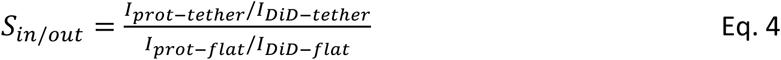

where 0 ≤ *S_in_*_/*out*_ < 1 indicates depletion and *S_in_*_/*out*_ > 1 represents sorting to the tether. The protein signals are normalized with the intensity *I_DiD_*_−*tet*ℎ*er*/*flat*_ of the curvature insensitive membrane marker DiD ^44^, to correct for out of focus effects and different tether sizes (Supplementary Figure S9).

The measured sorting indices for MUC5AC showed slight de-sorting from the in-tether and no effect for the out-tether (Fig. 3a), suggesting only a mild preference for flat surfaces at the tested expression levels. This agrees with the unchanged *C_0eff_* compared to wild-type HEK293 cells (Fig. 2c). In contrast, both expressed (cytosolic) Annexin A5 and its cognate protein Annexin A4 have been shown to sort into out-tethers in isolated GPMVs, a behaviour linked to curvature-inducing and -sensing capacity ^11,22,52^. Here we tested sorting of purified Annexin A5 extracellularly added to GPMV-cells. We found high sorting of Annexin A5 into in-tethers and depletion from out-tethers, consistent with its negative curvature-sensing ability, and in agreement with the opposite trend for cytosolic Annexin in previous works ^11,22^ (Fig. 3a). As expected, the sorting index decreased at high Annexin membrane coverage (Supplementary Figure S12a). Tracking the sorting index over the tether radius suggests a preferred curvature radius of ≈ 30 − 50 *nm* for Annexin A5 (Supplementary Figure S12b).

**Figure 3:**
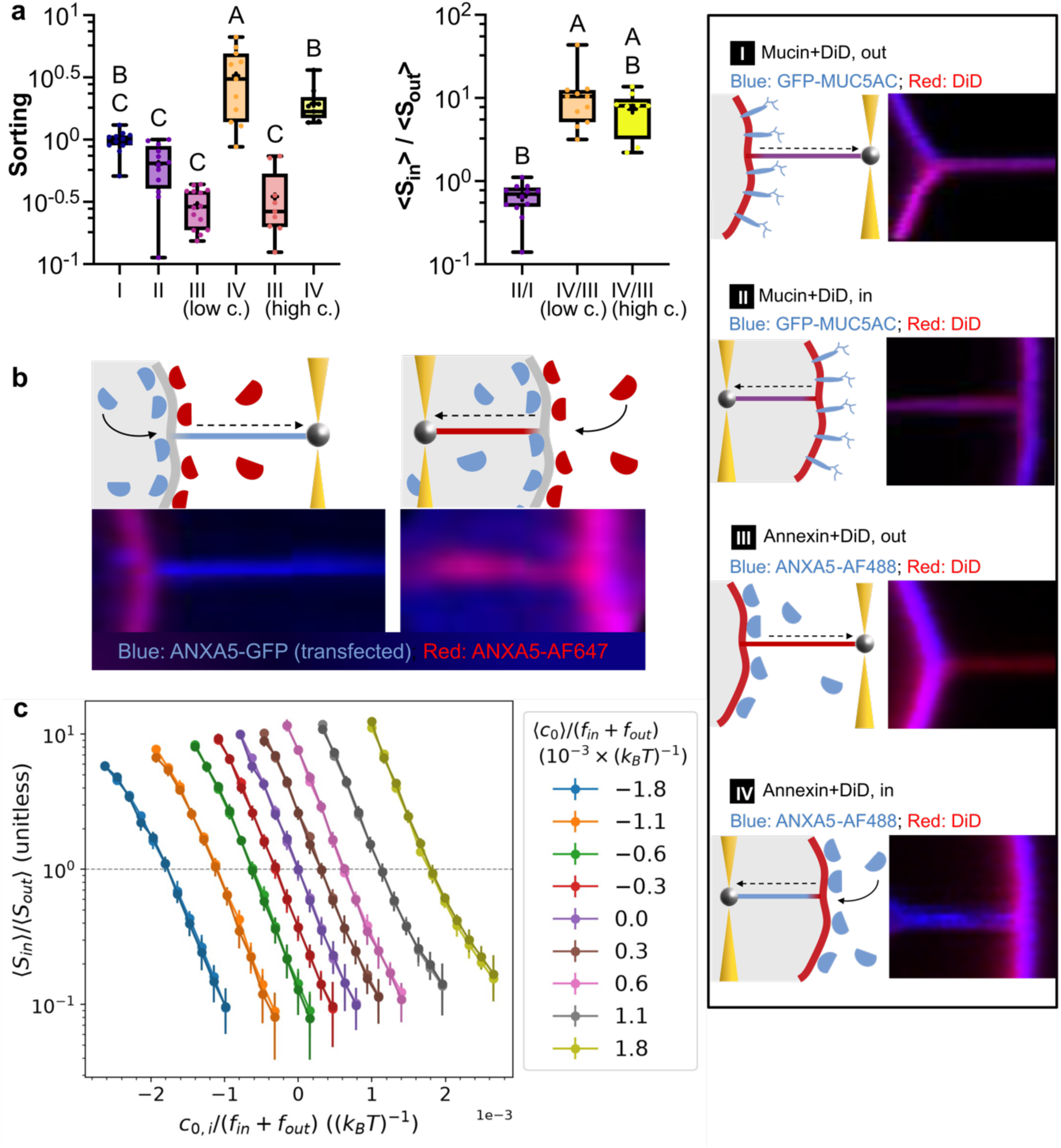
Curvature sorting of proteins. **(a)** In and out sorting on cell attached GPMVs expressing MUC5AC or exposed to two different concentrations (high c. and low c.) of extracellularly added Annexin A5-AF488, each showing different trends for curvature sorting. The right panel shows sorting ratios. The legend depicts schematics and confocal microscopy images of the conditions. Each box represents the interquartile range (IQR), with the median and mean (×) indicated. Groups with different capital letters differ significantly (P<0.05), whereas groups sharing the same letter do not (ordinary one-way ANOVA with Tukey’s multiple-comparisons test). Number of experimental days, n, and datapoints, N: n_mucin_ = 5, N_mucin_ = 12; n_low-C-Annexin_ = 5, N_low-C-Annexin_ = 11; n_high-C-Annexin_ = 3, N_high-C-Annexin_ = 9. **(b)** Confocal fluorescence images of a cell attached GPMV with an extended membrane tether, showing an overlay of expressed Annexin A5-GFP on the inside of the membrane (blue), and purified Annexin A5-AF647 outside (red). The expressed annexin from the inside sorts into the out-tether, and the purified Annexin A5-AF647 added to the outer leaflet sorts into the in-tether. **(c)** Sorting ratios in simulations of the different inclusion species as a function of their individual species curvature *c_0_*_,*i*_ scaled by the sum of in- and out-pulling forces to be independent of simulation units. Colors indicate the mean curvature ⟨*c_0_*⟩ of the curvature field/distribution. An overlay of the results for gaussian distributions and uniform distributions is shown, where the lighter colors mark data from uniform curvature distributions, the darker colors from gaussian curvature distributions. Each datapoint denotes the mean over frames from the last 1 × 10^7^ MC steps of 5 independent replicas, errorbars represent SEM over the 5 replicas.

We validated that Annexin A5 can sort to opposing curvatures when facing opposing leaflets at the single cell level, by extracellularly adding Annexin A5 probes labelled with AF647 to GPMV-cells that already express Annexin A5-GFP (Figure 3b). In this qualitatively instructive setup we can’t quantify sorting as done above, since there is no DiD membrane marker (due to spectral overlap with AF647), but it is possible to compare the ratios of both proteins in each tether. 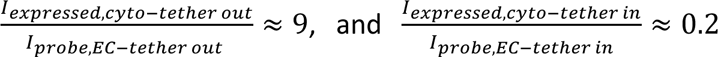 reveal a clear enrichment of the cytosolic expressed annexin in the out-tether, consistent with previous studies ^11,22,52^, and of the extracellular purified annexin in the in-tether with the corresponding de-sorting in the opposing direction for each.

### 3.2 Curvature-induced sorting in simulations and theory

Next, we compared the experimental sorting data to the simulations, building a relation between the mean spontaneous curvature ⟨*c_0_*⟩ of a heterogeneous membrane and the individual protein of interest sorting *S_in_*_/*out*_. In line with the experimental sorting index we defined a sorting ratio for an inclusion type *i* with spontaneous curvature *c_0_*_,*i*_ to an in- vs. an out-tether of the same system as 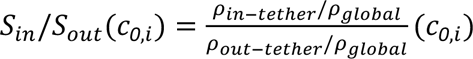. Analysis of simulated inclusion distributions (gaussian, uniform or asymmetric) showed sorting according to curvature preference, consistent for all types and widths of the distributions, indicating only a negligible influence of the distribution shape (Fig. 3c, Supplementary Figure S13). The sorting ratio of an inclusion species depended on the mean ⟨*c_0_*⟩ of the distribution that they were part of and was larger, the more negative the species’ curvature *c_0_*_,*i*_ is relative to all other curvatures *c_0_*_,j≠*i*_ in the same distribution.

To gain a better understanding, we described the system theoretically as two coupled compartments (tether and flat membrane) exchanging particles, treated within a grand-canonical ensemble with fixed total particle number (Supplementary Note 3). The membrane surface is divided into area elements of equal size *a_0_*, and assigned one inclusion to each element. A species’ theoretical sorting ratio is then:

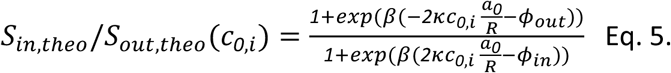

Here *R* is the tether radius (Eq. 2) and *φ_in_*_/*out*_ are Lagrange multipliers recovered through numerical solution of the boundary conditions (Supplementary Note 3, Eq. S3-8).

The theoretical sorting ratio is independent of the underlying distribution shape in an approximation to linear order, for all distributions with the same mean ⟨*c_0_*⟩ and curvatures significantly smaller than the tether radius *c_0_*_,*i*_*a_0_*/*R* << 1 (Supplementary Note 3, Eqs. S3-12 to S3-16). Deviations in the theoretical sorting ratio between different symmetric and asymmetric distributions are vanishingly small for the biologically relevant curvature ranges used in this work (see Supplementary Figure S14), in agreement with our simulation results. Hence, a priori knowledge of the experimental curvature distribution, such as the number of different species, is not required to compare experimental sorting ratios with simulated or theoretical sorting ratios.

Deviation between simulation and theoretical prediction of the sorting ratio occurred at larger |*c_0_*_,*i*_ | (Supplementary Fig. S15). Since the simulations allow changes to the tether radius due to the curvature preference of inclusions that sort to it, we confirmed that this deviation persists even when replacing the theory radius by *R_in_*, *R_out_* measured from simulations (Supplementary Figure S16). Therefore, these differences are most likely caused by the membrane fluctuations included in simulations ^53^, for whose influence on sorting the theory does not account.

### 3.3 Measuring individual protein curvature in cell membranes via sorting and experiment-simulation-pipeline

To extract the spontaneous curvature of a specific protein species in its native membrane environment from experiments we therefore propose using a simulated sorting diagram (Fig. 3c) with the following approach: (i) measure *Δf* to find the mean spontaneous curvature ⟨*c_0_*⟩ of the heterogeneous membrane containing the protein of interest, (ii) measure the sorting ratio of that protein via fluorescent labels (as in Fig. 3 a), (iii) read off *c_0_*_,*i*_ for the species of interest *i* on the corresponding ⟨*c_0_*⟩-curve from a simulated diagram like Fig. 3c and convert to physical units.

For Annexin A5, a previously determined spontaneous curvature from fully atomistic molecular dynamics simulations (aaMD) *c_0_*_,*Anx*_ = −0.047 ± 0.006 *nm*^−^*^1^*, see Ref. ^52^, exists as a benchmark. To test our concept, we determine the curvature of Annexin A5 from our dual-tether assay both at higher and lower concentrations (5*μL* and 50*μL* probes). The mean spontaneous curvature of annexin-decorated tethers for both cases is found as ⟨*c_0_*⟩/(*f_in_* + *f_out_*) = −(1.8 ± 0.3) × 10^−3^*k_B_T*^−1^ in Fig. 2f (see gridded version Supplementary Figure S17). Using the experimental sorting ratios (Table 1) and the exponential behaviour of *S_in_*/*S_out_* it is easy to read off *c_0_*_,*Anx*_/(*f_in_* + *f_out_*) on the logarithmic scale at this mean curvature (blue data in Fig. 3c, gridded version in Supplementary Figure S18). To convert this to explicit curvature values one can use the size of Annexin A5 from the aaMD reference (area occupied by one individual protein, see Ref. ^11,52^) and the average area of a simulation vertex (Table 1). This yields *c_0_*_,*Anx*_ *_5ul_* = −0.36 ± 0.02 *l_DTS_*^−^*^1^* = −0.056 ± 0.003 *nm*^−^*^1^* and *c_0_*_,*Anx*_ *_50ul_* = −0.32 ± 0.04 *l_DTS_*^−^*^1^*= −0.050 ± 0.006 *nm*^−^*^1^*. These curvatures measured via our pipeline are in very close agreement with the results determined through aaMD simulations.

**Table 1:** Measurement of Annexin A5 spontaneous curvature through sorting ratios on in- and out-tethers in combination with mesoscale simulations.

| ANXA5 probe volume | $S_{in}/S_{out}$ | $\frac{c_{0,Anx} l}{f_{in} + f_{out}}$ in $[10^{-3} k_B T^{-1}]$ | $\sqrt{\frac{A_{Anx}}{A_{v,sim}}}$ in $[\frac{nm}{l_{DTS}}]$ | $c_{0,Anx}$ in $[\frac{1}{nm}]$<br>converted via area under protein |
| --- | --- | --- | --- | --- |
| $5\mu\text{L}$ | $12 \pm 3$ | $-3.0 \pm 0.1$ | 6.4 | $-0.056 \pm 0.003$ |
| $50\mu\text{L}$ | $7 \pm 4$ | $-2.7 \pm 0.5$ | 6.4 | $-0.050 \pm 0.006$ |

Our approach opens a direct route to experimentally measure a protein curvature imprint in situ in native membranes, where the endogenous lipid composition, membrane asymmetry, and molecular interactions are naturally preserved, whereas aaMD relies on computationally intensive simulations that can only capture a highly limited subset of the native environment.

## DISCUSSION

Devising methods to reliably measure the spontaneous curvature of cellular plasma membrane that may express a complex mixture of endogenous curvature active proteins is highly important due to the implications of spontaneous curvature on a variety of biological processes. In this work, we introduced a measurement pipeline that combines dual-direction tether pulling from cell attached GPMVs using optical tweezers and mesoscale computer simulations to address this. We showed that the force difference from dual-direction tether pulling provides a robust measure of the mean of the heterogeneous spontaneous curvature distribution of cell-attached GPMVs. While the curvature sensed is not consistently identical to the mean ⟨*c_0_*⟩, it remains directly proportional and experimental force measurements can be easily translated to ⟨*c_0_*⟩ with the help of mesoscale simulations. Notably, our scheme can be applied on systems highly covered with various curvature-active protein types and deliver reliable estimates of the underlying ⟨*c_0_*⟩ independent of the specific protein and lipid distribution shape. Hence, a dual-direction tether pulling assay on cell-attached GPMVs opens a pathway to extend previously established methods beyond simple model systems like GUVs and to uncover the physical properties of various types of cell and organelle membranes and their membrane-protein populations which are at the core of cellular shape-function relations. Further studies on different cell types may enable a platform to compare their underlying distributions, with insights into their unique function.

A clear outcome of our analysis is that HEK293-derived GPMVs exhibit a negative effective spontaneous curvature of approximately *C_0eff_* = ⟨*c_0_*⟩ = −0.008 *nm*^−^*^1^*. Due to plasma membrane heterogeneity this value should be interpreted as an effective mean curvature preference of the native lipid and protein ensemble retained in the GPMV. One plausible contribution is transbilayer lipid abundance asymmetry. Recent work on human erythrocyte plasma membranes ^5^ showed that the cytoplasmic leaflet can contain over 50% more phospholipids than the exoplasmic leaflet, an imbalance enabled by asymmetric interleaflet cholesterol distribution and associated with leaflet-specific stresses. Although not directly quantitatively transferable from erythrocytes to HEK293 membranes, this provides a plausible physical framework in which excess effective area or altered tension in the cytoplasmic leaflet could favor inward bending and contribute to the negative spontaneous curvature observed here. Consistently, the systematic curvature shifts through sucrose and extracellular Annexin A5 underlined how leaflet-specific asymmetries and membrane-bound proteins tune the effective spontaneous curvature of plasma membranes.

Unlike peripherally acting sucrose or Annexin A5, the mucin reporter construct MUC5AC is an integral transmembrane component whose mechanical contribution reflects the combined properties of its extracellular domain, transmembrane anchor, and local membrane organization. The absence of a measurable curvature change with our MUC5AC does not allow general conclusions regarding membrane-associated mucins. However, future work could specifically compare different mucin reporters with other transmembrane proteins known for strong curvature-induction but previously not quantifiable in situ, such as the ion channels Piezo1 and KvAP ^54^. This would allow deeper insights on the differences in membrane curvature induction from transmembrane proteins versus peripheral agents.

We moreover introduced a curvature-sorting-based extension to the framework. The mean membrane spontaneous curvature ⟨*c_0_*⟩ is combined with additional measurements of the sorting ratio of a protein of interest and of inclusion mixtures in mesoscale simulations. Based on the example Annexin A5, we show how this enables measurement of the spontaneous curvature imprint of the protein of interest. One should note that high concentrations of the protein of interest may alter the bending rigidity of the membrane, which in turn influences the calculation of mean membrane spontaneous curvature ⟨*c_0_*⟩ and possibly also influences sorting behaviour. This could be mitigated in part by additional bending rigidity measurements to input the correct parameters for adjusted mesoscale simulation tables. Our approach may also be refined by including higher order interactions between inclusions that represent proteins. The dominating first order (local curvature induction) which we have considered here, already provided good compatibility with the experimental sorting data and all-atom molecular dynamics reference. However, as a second order correction, a local effect of inclusions on the bending rigidity of the membrane can lead to subtle clustering ^55^ and could additionally influence sorting. Including such advancements will require experimental data capable of capturing these small second order corrections. To generally broaden our assay to the influence of other protein-protein interactions on curvature sensing and generation, further experimental insights, e.g. through cross-linking mass spectrometry approaches ^56,57^ could be beneficial.

Integrating experiments and simulations in a joint framework like ours to better capture biological membrane system complexity is an approach increasingly employed in recent works ^11,58–60^. While experiments directly capture the native system, the simulations provide a controlled, physics-based environment to isolate individual parameters or access experimentally unavailable length- or timescales. The present work demonstrates specifically how simulations can establish the necessary connection between the experimental system and its idealized theoretical description, providing a route to study the membrane-shaping properties of curvature-active proteins and other molecular agents in physiologically relevant environments.

## METHODS

### Cell culture

A transmembrane GFP-tagged MUC5AC tandem repeat (TR) reporter was designed by fusing the human MUC1 signal peptide (amino acids 1-62, Uniprot P15921), a Flag-tag, EGFP (enhanced cyan fluorescent protein), the human MUC5AC TR sequence (amino acids 2708-2850, Uniprot P98088), and the membrane anchoring domain of human MUC1 (amino acids 1042–1138). The reporter was stably expressed in HEK293 wild-type (WT) under selection with G418 (500 ug/mL) as previously described (Ref. ^32^). HEK293 WT cells and stable MUC5AC reporter-expressing cells were cultured in Dulbecco’s Modified Eagle Medium (DMEM; Gibco, Cat. No. 392-0415) supplemented with 10% Fetal Bovine Serum (FBS, Sigma-Aldrich Cat. No. F7524) and 1% Penicillin-Streptomycin (Gibco, Cat. No. 15140122) in a humidified incubator at 37°C and 5% CO2. Cells were passaged at 70 − 90% confluency every 3-4 days and were used up to passage 20. For passaging, cells were washed once with 5 mL phosphate-buffered saline (PBS), incubated with 0.5 mL TrypLE Express for 5 min to detach the cells, collected by centrifugation, and resuspended in 5 mL of fresh medium. The concentration of suspended cells was determined in an Invitrogen Countess Automated Cell Counter. Approximately 450000 cells were then seeded in 5 ml of fresh media. Cell identity was authenticated by the vendor. Cells were routinely tested to ensure absence of mycoplasma.

### Sample preparation

The cell lines were plated using a 20-times diluted resuspension medium in 3.5 mm round glass bottom dishes (Mattek, Cat. No. P35G-1.5-10-C) in 2 mL of cell culture medium and incubated at 37°C in 5% CO_2_ overnight. Before cell plating, the glass dishes were coated with Poly-L-lysine (PLL, Sigma-Aldrich, Cat. No. P8920) by incubating with a 100 µg/mL solution in distilled water (Gibco, Cat. No. 12589069) for 30 minutes and then washed two times with excess distilled water.

After one day of incubation at 37 ◦C and 5% CO_2_, when required, cells at ∼50% confluency were transfected overnight with 2 µg Annexin A5-GFP in 1 mL Opti-MEM medium (Invitrogen, Cat. No. 11058021) using 2.5 µL of Lipofectamine LTX and 2 µL of plus reagent (Invitrogen, Cat. No. 15338030) as directed by the manufacturer’s protocol.

The next day, cells at ∼70% confluency were labeled in 1 ml of regular media with 2 µM DiD (Invitrogen, Cat. No. 10798263) and incubated at 37◦C and 5% CO_2_ for 15 minutes. Following labeling, the cells were washed twice with 2 ml of GPMVs buffer (150mM NaCl, 20mM HEPES, 2mM CaCl_2_, pH corrected to 7.4 using NaOH), and then 1 ml of the vesiculation buffer containing GPMVs buffer and 1mM N-Ethylmaleimide (NEM, Sigma-Aldrich, Cat. No. E3876-5G) was added. The cells were incubated for 1 hour at 37 ◦C and 5% CO_2_, and then 0.5μl of diluted 1.05 μm streptavidin-coated polystyrene beads (Spherotech, Cat. No. SVP-10-5) were added. When indicated, the 0.5M sucrose or either 5 µL (low concentration) or 50 µL (high concentration) Annexin A5 probes (fused to AF488, Invitrogen, Cat. No. A13201; or AF647, Invitrogen, Cat. No. A23204) are added at this step to 1 ml of vesiculation buffer. Finally, the chambers were sealed using a glass coverslip (#1.5H Glass, Ibidi, Cat. No. 10815) and the excess buffer was removed. The chambers were analyzed in the C-Trap (Lumicks) for up to 3 hours.

### Optical trapping and confocal imaging

Experiments were performed using the commercially available LUMICKS C-Trap optical tweezers system. A near-infrared trapping laser (1064 nm) was tightly focused through a Nikon 60× water-immersion objective (CFI Plan Apo, NA 1.2) to generate a highly stable gradient optical trap inside the experimental chamber.

A high-precision piezoelectric stage enabled accurate positioning of the trap to capture and move beads inside the microscopy chamber. Confocal fluorescence images were acquired simultaneously using two excitation lasers at 488 nm (for excitation of GFP-MUC5AC, Annexin A5-GFP, and Annexin A5-AF488 probe) and 640 nm (for excitation of DiD and Annexin A5-AF647 probe), with emission detected in two channels with a 512/25 nm filter, and a 680/42 nm filter, respectively. Pixel size was set to 100 nm, and the excitation laser powers were 3% (1.12 μW) for the 488 nm laser and 1% (0.94 μW) for the 640 nm laser.

For force detection, forward-scattered light from the trapping laser was collected by an oil-immersion condenser and directed onto a position-sensitive detector (PSD). The PSD recorded bead displacements at a 70 kHz sampling rate, allowing precise quantification of force changes during the experiments. For force measurements, in addition to confocal imaging, bright-field imaging of the sample plane was performed in parallel at 60 frames per second.

### Dual tether assay on cell attached GPMVs

Suitable GPMVs were approximately spherical, appeared intact and sufficiently separated from neighboring cell structures, were still firmly attached to their parent cell and mechanically stable during stage movement.

Force application was initiated by moving the stage to bring the optically trapped bead into contact with the GPMV. After contact was established, the bead was carefully pushed towards the GPMV surface (∼5 s, approximately 20 pN final contact repulsive force) to enable firm attachment of the bead via non-specific interactions. After ∼30 s rest at the bead-GPMV contact area, an automated 0.5 μm/s stage movement was initiated in a direction perpendicular to the GPMV surface: first away from the cell, generating an out-tether (positive curvature) which is held in focus for at least 1 minute; and then towards the cell, generating an in-tether (negative curvature) which is held in focus for at least 1 minute.

The force required to hold a stable tether was quantified using a 2% calibrated laser power, corresponding to 44 mW. A low laser power provided a more stable force measurement and minimized the chance of trapping other particles present in the chamber (e.g., other beads or vesicles). However, before the active nanostage moving period (tether extension periods) the laser power was increased from 2% to 15%, corresponding to 380 mW. Increasing the laser power re-positioned the bead within the trap center and ensured that the bead stayed in the trap during the tether formation regime. The laser power is decreased back to 2% once the nanostage movement stops, corresponding to the moment the tether length equals at least 5µ*m* for the out-tether, or 3µ*m* for the in-tether.

Once the stable tether is formed, the stage z position is changed in 100 nm steps until the best tether focus is found. The sorting and force analysis are done exclusively in regions where the tether was in focus, determined by manual inspection of the best in-focus frame for each tether confocal image, and using a force vs time plot, where the corresponding confocal scan start-end frames are indicated.

When holding in tethers, there is a probability that the tether adopts a different orientation in the z-direction to minimize its length, and therefore goes out of the trap focus and the confocal image plane, which consequently lead to inaccurate force measurements and imaging (Supplementary Figure S1). This is typically corrected by bringing the trap closer to the original GPMV surface entrance point and optimizing the stage z-position. This effect limits the experimentally achievable in-tether length.

### Force quantification

For quantitative force measurements, the trap position was held stationary, and the sample was moved using a stage controller. A single bead was optically trapped initially at 2% (44 mW) laser power and calibration were achieved by automated fitting of a Lorentzian function to the power spectral density of the Brownian bead fluctuations (average spring constant 0.15 ± 0.07pN/nm, N = 68). The base-line force was set to zero corresponding to a position of the bead at the center of the trap.

Force magnitude plateaus for the out- or in-tether were manually selected from a force vs time profile, and approximately 20 seconds were averaged at the regions where the tether was in-focus. The corresponding averages were used in equation 1 in the form *Δf* = *f_in_* − *f_out_* and equation (S1-6).

Real-time force data and confocal images were exported from Bluelake HDF5 files and analyzed using custom scripts in the Pylake (v1.8.0, LUMICKS) Python (version 3.12) package. Force signals were down sampled to 781.25 Hz for analysis and to 15 Hz for plotting.

### Analysis of Protein Sorting to Tethers pulled from GPMVs

The vesicle shape was manually approximated by fitting a circle (Supplementary Figure S19a, green), from which a 180° arc (upper hemisphere – vesicle marker) was selected with an arc width of 3.1 μm, centered at the fitted radius. A rectangular crop of width 3.1 μm (Supplementary Figure S19a, green rectangle-tube marker), centered along the line connecting two manually selected points at the tether ends, was also created for further analysis. The intensity of protein-and membrane dyes were then measured within these regions to calculate the sorting as 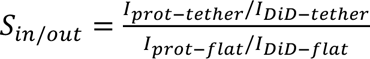 (see Eq. 4 above). The intensity of protein dyes was associated with protein surface density. The intensity of membrane dyes was used as a relative tether size and for size comparison of the in-tether and out-tether. For more details on how the relevant intensity is quantified, see Supplementary Figure S9.

### Simulation Methods

Mesoscale simulations were performed using Dynamically Triangulated Surfaces (DTS) with the software FreeDTS ^20^ which has shown to reproduce theoretically tractable thermodynamical behaviours of membranes such as the undulation spectrum ^27,61^. DTS simulations generally are a highly effective approach to simulating biological membranes at large scales ^62,63^. There are a few different models and software packages available for performing mesoscale simulations ^64–70^. Descriptions of the FreeDTS simulations framework and further references can be found in Ref. ^20,59^ but for the benefit of the reader, in the following paragraphs we revisit the aspects which are relevant to the work presented here. Within the framework of FreeDTS, a membrane is modelled as a triangulated surface of *N_v_* vertices, *N_e_* edges and *N_T_* triangles, with a characteristic length unit *l_DTS_* being the minimal distance between any two vertices of the membrane mesh. The curvature at each vertex is calculated via the shape operator formulation ^27^. In order to obtain the bending energy of the surface, a set of discretised geometrical operations is applied at each vertex to determine its unit normal *N*^b^*_u_*, surface area *A_u_*, principal curvatures (*c*_1*u*_, *c*_2*u*_) and principal directions (*X*_1_(*υ*), *X*_2_(*υ*)). These yield the bending energy of the surface (*E_B_*) described by a discretised form of the Helfrich Hamiltonian:

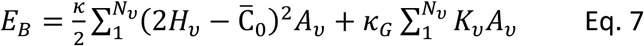

in terms of the mean 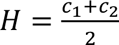, and Gaussian curvature, *K* = *c*_1_*c*_2_ and with the bending modulus *κ*, Gaussian modulus *κ_G_*, and spontaneous curvature C^j^_0_. The latter represents a possible asymmetry between the two monolayers (C^j^_0_ = 0 for a symmetric membrane). The equilibrium configurations of the system are sampled using a Monte Carlo (MC) metropolis algorithm, consisting of multiple moves i.e., vertex position update, edge flip and Kawasaki moves to change the position of the inclusions (see Ref. ^20^ for more details). A MC step corresponds to *N_v_* attempts to move a vertex and *N_e_* attempts to flip links and *N_i_* Kawasaki moves to attempt moving inclusions to new positions ^71^.

For constant frame tension simulations in PBC a position rescaling algorithm adjusts the box size ^27^ to balance the internal and the external stresses. For every X^th^ Monte Carlo (MC) step, an additional box change attempt is made, with these moves also being subject to a metropolis criterion for acceptance or rejection. The algorithm couples the system energy to

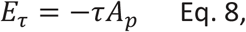

where *A_p_* is the projected area which can vary and adapt, corresponding to the area of the periodic box in x- and y-direction. Once the simulation is performed and the system reaches a minimal free energy, the average internal stresses from the internal energetic terms, e.g. from stretching and bending, balance those externally imposed by mechanical tension. For details on the tension algorithm please see Ref. ^27,59^.

### Generating configurations for tethers on flat membrane patches with PBC

To obtain a membrane tether in PBC, a flat membrane configuration was first relaxed and equilibrated in PBC with a constant tension of *τ* = 0*k_B_T*, allowing for adjustment of the box size. We then subjected our membrane to lateral tension with parameter *τ* > 0 (in the x-y-plane) and slowly pulled a membrane tether to desired length 20 ≤ *L* ≤ 120 in the center of mass (COM) of the membrane, see Supplementary Figure S19b. The pulling was realised by applying a harmonic potential along the z-direction between one vertex located close to the COM of the flat membrane in the periodic box at position 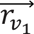 (envisioned tip of the tube) and all other vertices of the membrane mesh with centre of geometry 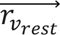 :

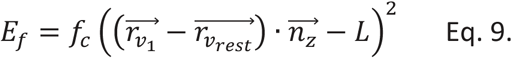

Here, *f_c_* denotes the force constant, and *L* the target distance between the tip of the tube and the center of geometry of all other vertices, which provides a measure of the target tube length (please note that *L* it is not a measure of the full final tube length, but the distance between the tube tip and the COM of all other vertices). In order to reach a well equilibrated tether at all lengths, the pulling was performed by starting with a small *L* = 10*l_DTS_*, which was increased by 0.05*l_DTS_* every 2000 MC steps until the desired *L* was reached.

These tether configurations were used as the basis to set up any further simulations to evaluate the force elongation curve, the effect of curvature inducing proteins and any other additional conditions. That means that all results provided in this work were obtained from taking a pre-equilibrated tether of the desired certain length, and then additionally equilibrating this respective configuration under the newly appended constraints or conditions, e.g. lower tension *τ*, for 8 − 12 × 10^7^ MC steps. Of those, configurations of the last 1 − 2 × 10^7^ MC steps were used for analysis.

### Converting simulation results to physical units

To compare force measurement simulations to experimental measurements, we can convert the internal simulation lengthscale *l_DTS_* to physical units by choosing the membrane tension as a reference. From experiments, tension is not directly available as an independent measure, but can be calculated from force measurements, using the same bending rigidity *κ* = 82.32 *pNnm* = 20*k_B_T* as in simulations. For the HEK-cell controls we find *τ_HEK293_* ≈ *0*.*0197* ± *0*.*0029 pN*/*nm*, while for simulations we use *τ_sim_* = 2*k_B_T*/*l_DTS_*^2^. Then *l_DTS_* ≈ 20.4 ± 1.5 *nm*. Often the conversion can be avoided, by reporting quantities that are available directly both to experiment and simulation. To do so, we report both force or force difference measurements as well as curvatures rescaled by the sum of in-pulling and out-pulling force (*f_in_* + *f_out_*), i.e. *Δf*/(*f_in_* + *f_out_*) and ⟨*c_0_*⟩/(*f_in_* + *f_out_*) as done for example in Fig. 2 and Fig. 3. Since the forces are directly measured both in experiments and simulations, this rescaling is independent of assumptions about the bending rigidity in experiments (within biologically relevant scales), and provides rescaled forces unitless, and rescaled curvatures in units of 1/energy. Lastly, to compare sorting measurements of a protein of interest, one can furthermore equate the protein size (area occupied by the protein on a membrane surface) and the average vertex area *A_u_* from simulations to correctly convert *l_DTS_* to physical units for curvatures.

### Analysis

Every simulation was run in PBC with 5 replicas, except for the force elongation curve (Supplementary Figure S3) where 13 replicas were used. Each datapoint presented in this work for pulling force or tether radius hence corresponds to the mean of the values recovered for these replicas. Errorbars indicate the standard deviation.

The pulling force is measured in simulations as the derivative of eq. (9) with respect to the 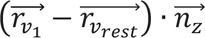, i.e.

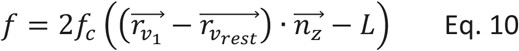

and can be read from the energy output files of the simulations of interest, where it is saved every 100 MC steps. The tether radius can be determined from the simulated topology through knowledge of the vertex sitting at the tether tip (see also Supplementary Figure S19c). Using its position, we consider only those vertices of the topology file, which lie between *z_tip_* − *0*.*9l_0_* and *z_tip_* − *0*.*05l_0_* along the tube with length *l_0_*and target length *L* for the harmonic potential, avoiding the tube tip itself as well as the connective region between tether and flat part of the membrane. The chosen vertices are divided into slices of *dz* = 2 *l_DTS_* along the z-axis, as the tether is pulled in parallel to it. We then fit a circle to the positions of vertices within each slice (*x_i_*, *y_i_*)*_i_*_∈{*vertices*_ *_in slice_*_}_ to approximate the radius*r_slice_* of the tether per slice. The overall radius of the tether is calculated as the mean of the slice radii. This approach ensures that possible tilting of the tether (i.e. deviations of its axis from the global z-axis) as well as radius fluctuations along the tether length are accounted for. To analyse sorting of inclusions the membrane is divided into the tether and flat region at *z_cutoff_* = *min*(*z*) + 6*l_DTS_*. Any inclusion positioned higher is counted as situated on the tether, any lower as situated on the flat region. The vertices are sorted into two groups in the same way and the area of each region is calculated as the sum of its vertex areas. This allows calculation of density for each inclusion species in each region and each frame of the simulation. We analyse 5 replica of each simulation and report sorting measures as the mean over these.

### Heterogeneous membranes

To generate a heterogeneous membrane, we used the concept of inclusions in FreeDTS, an object that sits on vertices so that there is at most one inclusion per membrane vertex ^12^. To study the effect of curvature inducing proteins on membrane tethers, we chose inclusions with symmetric interactions with the membrane (inclusion type 1 in FreeDTS):

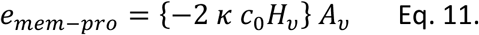

Here, we use a simple model without local change to the membrane bending rigidity from to the protein, and *c*_0_ is the parameter for the local curvature preference of that protein. For more details see ^20^. Membranes were decorated by ≈100% of inclusions of multiple types with a normal, uniform or asymmetric distribution (i.e. the *c_0_* parameter was varied and the percentage of inclusions with a certain value of it was chosen according to the distribution around a chosen mean and standard deviation). No direct protein-protein interactions were employed.

## Acknowledgements

The research of BG and WP was supported by the Novo Nordisk Foundation grants NNF18SA0035142 and NNF22OC0079182 and Independent Research Fund Denmark grant No. 10.46540/2064-00032B. LHA and PMB acknowledge support from the Novo Nordisk Foundation grants NNF23OC008432, NNF20OC0061176 and NFF25OC0099835.

## Author Contributions

WP and PM conceived the original idea. BG prepared the initial draft of the manuscript, which was subsequently revised and developed by BG, LHA, WP, and PM. LHA performed experiments and analysed them, with contributions from ED and NK. VTR, YB and GMP were involved in development of the experimental assay. JN, YN, KTS, and HJ provided experimental expertise related to annexin- and mucin-expressing cell lines. BG performed simulations, obtained the results, developed the theory and partially analysed experimental data. All authors discussed the results, commented on the manuscript throughout its preparation, and approved the final version.

## Data Availability

All relevant files related to this manuscript are available from the corresponding author upon reasonable request. Source data will be provided with this paper. Simulation files and code used for analysis of simulations in this work will be deposited publicly available on Github.

## Code Availability

All relevant code related to this manuscript is freely available on Github: https://github.com/weria-pezeshkian/FreeDTS.

## BIBLIOGRAPHY

(1) McMahon, H. T.; Gallop, J. L. Membrane Curvature and Mechanisms of Dynamic Cell Membrane Remodelling. Nature 2005, 438 (7068), 590–596. 10.1038/nature04396.

(2) Zimmerberg, J.; Kozlov, M. M. How Proteins Produce Cellular Membrane Curvature. Nat. Rev. Mol. Cell Biol. 2006, 7 (1), 9–19. 10.1038/nrm1784.

(3) Bassereau, P.; Jin, R.; Baumgart, T.; Deserno, M.; Dimova, R.; Frolov, V. A.; Bashkirov, P. V.; Grubmüller, H.; Jahn, R.; Risselada, H. J.; Johannes, L.; Kozlov, M. M.; Lipowsky, R.; Pucadyil, T. J.; Zeno, W. F.; Stachowiak, J. C.; Stamou, D.; Breuer, A.; Lauritsen, L.; Simon, C.; Sykes, C.; Voth, G. A.; Weikl, T. R. The 2018 Biomembrane Curvature and Remodeling Roadmap. J. Phys. Appl. Phys. 2018, 51 (34), 343001. 10.1088/1361-6463/aacb98.

(4) Deserno, M. Biomembranes Balance Many Types of Leaflet Asymmetries. Curr. Opin. Struct. Biol. 2024, 87, 102832. 10.1016/j.sbi.2024.102832.

(5) Doktorova, M.; Symons, J. L.; Zhang, X.; Wang, H.-Y.; Schlegel, J.; Lorent, J. H.; Heberle, F. A.; Sezgin, E.; Lyman, E.; Levental, K. R.; Levental, I. Cell Membranes Sustain Phospholipid Imbalance via Cholesterol Asymmetry. Cell 2025, 188 (10), 2586–2602.e24. 10.1016/j.cell.2025.02.034.

(6) Structural and Functional Asymmetry of Plasma Membranes: Faraday Discussion 259; Royal Society of Chemistry, 2025.

(7) Hossein, A.; Deserno, M. Spontaneous Curvature, Differential Stress, and Bending Modulus of Asymmetric Lipid Membranes. Biophys. J. 2020, 118 (3), 624–642. 10.1016/j.bpj.2019.11.3398.

(8) Seifert, U. Configurations of Fluid Membranes and Vesicles. Adv. Phys. 1997, 46 (1), 13–137. 10.1080/00018739700101488.

(9) Seifert, U.; Berndl, K.; Lipowsky, R. Shape Transformations of Vesicles: Phase Diagram for Spontaneous-Curvature and Bilayer-Coupling Models. Phys. Rev. A 1991, 44 (2), 1182–1202. 10.1103/PhysRevA.44.1182.

(10) Johannes, L.; Parton, R. G.; Bassereau, P.; Mayor, S. Building Endocytic Pits without Clathrin. Nat. Rev. Mol. Cell Biol. 2015, 16 (5), 311–321. 10.1038/nrm3968.

(11) Dam Florentsen, C.; Kamp-Sonne, A.; Moreno-Pescador, G.; Pezeshkian, W.; Zanjani, A. A. H.; Khandelia, H.; Nylandsted, J.; Martin Bendix, P. Annexin A4 Trimers Are Recruited by High Membrane Curvatures in Giant Plasma Membrane Vesicles. Soft Matter 2021, 17 (2), 308–318. 10.1039/D0SM00241K.

(12) Lizarbe, M. A.; Barrasa, J. I.; Olmo, N.; Gavilanes, F.; Turnay, J. Annexin-Phospholipid Interactions. Functional Implications. Int. J. Mol. Sci. 2013, 14 (2), 2652–2683. 10.3390/ijms14022652.

(13) Pozrikidis, C. Resting Shape and Spontaneous Membrane Curvature of Red Blood Cells. Math. Med. Biol. J. IMA 2005, 22 (1), 34–52. 10.1093/imammb/dqh021.

(14) Jani, P.; Colville, M. J.; Park, S.; Ha, Y.; Paszek, M. J.; Abbott, N. L. Influence of the Glycocalyx on the Size and Mechanical Properties of Plasma Membrane-Derived Vesicles. Soft Matter 2025, 21 (3), 463–475. 10.1039/D4SM01317D.

(15) Heinrich, M. C.; Capraro, B. R.; Tian, A.; Isas, J. M.; Langen, R.; Baumgart, T. Quantifying Membrane Curvature Generation of Drosophila Amphiphysin N-BAR Domains. J. Phys. Chem. Lett. 2010, 1 (23), 3401–3406. 10.1021/jz101403q.

(16) Sorre, B.; Callan-Jones, A.; Manzi, J.; Goud, B.; Prost, J.; Bassereau, P.; Roux, A. Nature of Curvature Coupling of Amphiphysin with Membranes Depends on Its Bound Density. Proc. Natl. Acad. Sci. 2012, 109 (1), 173–178. 10.1073/pnas.1103594108.

(17) Karimi, M.; Steinkühler, J.; Roy, D.; Dasgupta, R.; Lipowsky, R.; Dimova, R. Asymmetric Ionic Conditions Generate Large Membrane Curvatures. Nano Lett. 2018, 18 (12), 7816–7821. 10.1021/acs.nanolett.8b03584.

(18) Dasgupta, R.; Miettinen, M. S.; Fricke, N.; Lipowsky, R.; Dimova, R. The Glycolipid GM1 Reshapes Asymmetric Biomembranes and Giant Vesicles by Curvature Generation. Proc. Natl. Acad. Sci. 2018, 115 (22), 5756–5761. 10.1073/pnas.1722320115.

(19) Barbetta, C.; Fournier, J.-B. On the Fluctuations of the Force Exerted by a Lipid Nanotubule. Eur. Phys. J. E 2009, 29 (2), 183–189. 10.1140/epje/i2009-10468-8.

(20) Pezeshkian, W.; Ipsen, J. H. Mesoscale Simulation of Biomembranes with FreeDTS. Nat. Commun. 2024, 15 (1), 548. 10.1038/s41467-024-44819-w.

(21) Steinkühler, J.; Sezgin, E.; Urbančič, I.; Eggeling, C.; Dimova, R. Mechanical Properties of Plasma Membrane Vesicles Correlate with Lipid Order, Viscosity and Cell Density. Commun. Biol. 2019, 2 (1), 337. 10.1038/s42003-019-0583-3.

(22) Moreno-Pescador, G.; Florentsen, C. D.; Østbye, H.; Sønder, S. L.; Boye, T. L.; Veje, E. L.; Sonne, A. K.; Semsey, S.; Nylandsted, J.; Daniels, R.; Bendix, P. M. Curvature- and Phase-Induced Protein Sorting Quantified in Transfected Cell-Derived Giant Vesicles. ACS Nano 2019, 13 (6), 6689–6701. 10.1021/acsnano.9b01052.

(23) Doktorova, M.; Harries, D.; Khelashvili, G. Determination of Bending Rigidity and Tilt Modulus of Lipid Membranes from Real-Space Fluctuation Analysis of Molecular Dynamics Simulations. Phys. Chem. Chem. Phys. PCCP 2017, 19 (25), 16806–16818. 10.1039/c7cp01921a.

(24) Harmandaris, V. A.; Deserno, M. A Novel Method for Measuring the Bending Rigidity of Model Lipid Membranes by Simulating Tethers. J. Chem. Phys. 2006, 125 (20), 204905. 10.1063/1.2372761.

(25) Hu, M. (胡明暘); Diggins, P., IV; Deserno, M. Determining the Bending Modulus of a Lipid Membrane by Simulating Buckling. J. Chem. Phys. 2013, 138 (21), 214110. 10.1063/1.4808077.

(26) Eid, J.; Razmazma, H.; Jraij, A.; Ebrahimi, A.; Monticelli, L. On Calculating the Bending Modulus of Lipid Bilayer Membranes from Buckling Simulations. J. Phys. Chem. B 2020, 124 (29), 6299–6311. 10.1021/acs.jpcb.0c04253.

(27) Pezeshkian, W.; Ipsen, J. H. Fluctuations and Conformational Stability of a Membrane Patch with Curvature Inducing Inclusions. Soft Matter 2019, 15 (48), 9974–9981. 10.1039/C9SM01762C.

(28) Lipowsky, R. The Many Faces of Membrane Tension for Biomembranes and Vesicles. Faraday Discuss. 2025, 259 (0), 234–263. 10.1039/D4FD00184B.

(29) Raucher, D.; Sheetz, M. P. Characteristics of a Membrane Reservoir Buffering Membrane Tension. Biophys. J. 1999, 77 (4), 1992–2002. 10.1016/S0006-3495(99)77040-2.

(30) Muñoz-Basagoiti, M.; Frey, F.; Meadowcroft, B.; Amaral, M.; Prada, A.; Šarić, A. A Tutorial for Mesoscale Computer Simulations of Lipid Membranes: Tether Pulling, Tubulation and Fluctuations. Soft Matter 2025. 10.1039/D5SM00148J.

(31) Shurer, C. R.; Kuo, J. C.-H.; Roberts, L. M.; Gandhi, J. G.; Colville, M. J.; Enoki, T. A.; Pan, H.; Su, J.; Noble, J. M.; Hollander, M. J.; O’Donnell, J. P.; Yin, R.; Pedram, K.; Möckl, L.; Kourkoutis, L. F.; Moerner, W. E.; Bertozzi, C. R.; Feigenson, G. W.; Reesink, H. L.; Paszek, M. J. Physical Principles of Membrane Shape Regulation by the Glycocalyx. Cell 2019, 177 (7), 1757–1770.e21. 10.1016/j.cell.2019.04.017.

(32) Nason, R.; Büll, C.; Konstantinidi, A.; Sun, L.; Ye, Z.; Halim, A.; Du, W.; Sørensen, D. M.; Durbesson, F.; Furukawa, S.; Mandel, U.; Joshi, H. J.; Dworkin, L. A.; Hansen, L.; David, L.; Iverson, T. M.; Bensing, B. A.; Sullam, P. M.; Varki, A.; Vries, E. de; de Haan, C. A. M.; Vincentelli, R.; Henrissat, B.; Vakhrushev, S. Y.; Clausen, H.; Narimatsu, Y. Display of the Human Mucinome with Defined O-Glycans by Gene Engineered Cells. Nat. Commun. 2021, 12, 4070. 10.1038/s41467-021-24366-4.

(33) Bhatia, T.; Christ, S.; Steinkühler, J.; Dimova, R.; Lipowsky, R. Simple Sugars Shape Giant Vesicles into Multispheres with Many Membrane Necks. Soft Matter 2020, 16 (5), 1246–1258. 10.1039/C9SM01890E.

(34) Sens, P.; Plastino, J. Membrane Tension and Cytoskeleton Organization in Cell Motility. J. Phys. Condens. Matter 2015, 27 (27), 273103. 10.1088/0953-8984/27/27/273103.

(35) Gauthier, N. C.; Masters, T. A.; Sheetz, M. P. Mechanical Feedback between Membrane Tension and Dynamics. Trends Cell Biol. 2012, 22 (10), 527–535. 10.1016/j.tcb.2012.07.005.

(36) Hochmuth, R. M.; Marcus, W. D. Membrane Tethers Formed from Blood Cells with Available Area and Determination of Their Adhesion Energy. Biophys. J. 2002, 82 (6), 2964–2969. 10.1016/S0006-3495(02)75637-3.

(37) Evans, E.; Heinrich, V.; Ludwig, F.; Rawicz, W. Dynamic Tension Spectroscopy and Strength of Biomembranes. Biophys. J. 2003, 85 (4), 2342–2350. 10.1016/S0006-3495(03)74658-X.

(38) Evans, E.; Smith, B. A. Kinetics of Hole Nucleation in Biomembrane Rupture. New J. Phys. 2011, 13 (9), 095010. 10.1088/1367-2630/13/9/095010.

(39) Portet, T.; Dimova, R. A New Method for Measuring Edge Tensions and Stability of Lipid Bilayers: Effect of Membrane Composition. Biophys. J. 2010, 99 (10), 3264–3273. 10.1016/j.bpj.2010.09.032.

(40) Hategan, A.; Law, R.; Kahn, S.; Discher, D. E. Adhesively-Tensed Cell Membranes: Lysis Kinetics and Atomic Force Microscopy Probing. Biophys. J. 2003, 85 (4), 2746–2759. 10.1016/S0006-3495(03)74697-9.

(41) Tsai, F.-C.; Bertin, A.; Bousquet, H.; Manzi, J.; Senju, Y.; Tsai, M.-C.; Picas, L.; Miserey-Lenkei, S.; Lappalainen, P.; Lemichez, E.; Coudrier, E.; Bassereau, P. Ezrin Enrichment on Curved Membranes Requires a Specific Conformation or Interaction with a Curvature-Sensitive Partner. eLife 2018, 7, e37262. 10.7554/eLife.37262.

(42) Pipathsouk, A.; Brunetti, R. M.; Town, J. P.; Graziano, B. R.; Breuer, A.; Pellett, P. A.; Marchuk, K.; Tran, N.-H. T.; Krummel, M. F.; Stamou, D.; Weiner, O. D. The WAVE Complex Associates with Sites of Saddle Membrane Curvature. J. Cell Biol. 2021, 220 (8), e202003086. 10.1083/jcb.202003086.

(43) Tsai, F.-C.; Henderson, J. M.; Jarin, Z.; Kremneva, E.; Senju, Y.; Pernier, J.; Mikhajlov, O.; Manzi, J.; Kogan, K.; Le Clainche, C.; Voth, G. A.; Lappalainen, P.; Bassereau, P. Activated I-BAR IRSp53 Clustering Controls the Formation of VASP-Actin–Based Membrane Protrusions. Sci. Adv. 2022, 8 (41), eabp8677. 10.1126/sciadv.abp8677.

(44) Larsen, J. B.; Rosholm, K. R.; Kennard, C.; Pedersen, S. L.; Munch, H. K.; Tkach, V.; Sakon, J. J.; Bjørnholm, T.; Weninger, K. R.; Bendix, P. M.; Jensen, K. J.; Hatzakis, N. S.; Uline, M. J.; Stamou, D. How Membrane Geometry Regulates Protein Sorting Independently of Mean Curvature. ACS Cent. Sci. 2020, 6 (7), 1159–1168. 10.1021/acscentsci.0c00419.

(45) Breuer, A.; Lauritsen, L.; Bertseva, E.; Vonkova, I.; Stamou, D. Quantitative Investigation of Negative Membrane Curvature Sensing and Generation by I-BARs in Filopodia of Living Cells. Soft Matter 2019, 15 (48), 9829–9839. 10.1039/C9SM01185D.

(46) Cohen, M. J.; Chirico, W. J.; Lipke, P. N. Through the Back Door: Unconventional Protein Secretion. Cell Surf. 2020, 6, 100045. 10.1016/j.tcsw.2020.100045.

(47) Popa, S. J.; Stewart, S. E.; Moreau, K. Unconventional Secretion of Annexins and Galectins. Semin. Cell Dev. Biol. 2018, 83, 42–50. 10.1016/j.semcdb.2018.02.022.

(48) Zhao, W.; Hanson, L.; Lou, H.-Y.; Akamatsu, M.; Chowdary, P. D.; Santoro, F.; Marks, J. R.; Grassart, A.; Drubin, D. G.; Cui, Y.; Cui, B. Nanoscale Manipulation of Membrane Curvature for Probing Endocytosis in Live Cells. Nat. Nanotechnol. 2017, 12 (8), 750–756. 10.1038/nnano.2017.98.

(49) Prévost, C.; Zhao, H.; Manzi, J.; Lemichez, E.; Lappalainen, P.; Callan-Jones, A.; Bassereau, P. IRSp53 Senses Negative Membrane Curvature and Phase Separates along Membrane Tubules. Nat. Commun. 2015, 6 (1), 8529. 10.1038/ncomms9529.

(50) Zhu, C.; Das, S. L.; Baumgart, T. Nonlinear Sorting, Curvature Generation, and Crowding of Endophilin N-BAR on Tubular Membranes. Biophys. J. 2012, 102 (8), 1837–1845. 10.1016/j.bpj.2012.03.039.

(51) Ramesh, P.; Baroji, Y. F.; Reihani, S. N. S.; Stamou, D.; Oddershede, L. B.; Bendix, P. M. FBAR Syndapin 1 Recognizes and Stabilizes Highly Curved Tubular Membranes in a Concentration Dependent Manner. Sci. Rep. 2013, 3, 1565. 10.1038/srep01565.

(52) Pandey, M. P.; Souza, P. C. T. de; Pezeshkian, W.; Khandelia, H. Bending of a Lipid Membrane Edge by Annexin A5 Trimers. Biophys. J. 2024, 123 (8), 1006–1014. 10.1016/j.bpj.2024.03.019.

(53) Andreghetti, D.; Dall’Asta, L.; Gamba, A.; Kolokolov, I.; Lebedev, V. Molecular Sorting on a Fluctuating Membrane. 2025.

(54) Cino, E. A.; Tieleman, D. P. Curvature Footprints of Transmembrane Proteins in Simulations with the Martini Force Field. J. Phys. Chem. B 2024, 128 (25), 5987–5994. 10.1021/acs.jpcb.4c01385.

(55) Vidal, A. B.; Pezeshkian, W. Multi-Body Fluctuation-Induced Forces Between Membrane Proteins: Insights from Mesoscale Simulations. bioRxiv September 17, 2025, p 2025.09.12.675822. 10.1101/2025.09.12.675822.

(56) Schuhmann, F.; Akkaya, K. C.; Puchkov, D.; Hohensee, S.; Lehmann, M.; Liu, F.; Pezeshkian, W. Integrative Molecular Dynamics Simulations Untangle Cross-Linking Data to Unveil Mitochondrial Protein Distributions. Angew. Chem. Int. Ed. 2025, 64 (6), e202417804. 10.1002/anie.202417804.

(57) Seggern, B. von; Sadeghi, M. Methods for Inferring Interaction Potentials from Cross-Linking Mass Spectrometry Data. arXiv June 4, 2026. 10.48550/arXiv.2606.05541.

(58) Argudo, D.; Bethel, N. P.; Marcoline, F. V.; Grabe, M. Continuum Descriptions of Membranes and Their Interaction with Proteins: Towards Chemically Accurate Models. Biochim. Biophys. Acta 2016, 1858 (7 Pt B), 1619–1634. 10.1016/j.bbamem.2016.02.003.

(59) Geiger, B. J.; Pezeshkian, W. Bimodal Mechanical Response of Membrane Necks: Implications for the Nuclear Envelope. ACS Nano 2025. 10.1021/acsnano.5c05817.

(60) Attar, A. G.; Paturej, J.; Sarýyer, O. S.; Banigan, E. J.; Erbaþ, A. Peripheral Heterochromatin Tethering Is Required for Chromatin-Based Nuclear Mechanical Response. Nucleic Acids Res. 2025, 53 (15), gkaf763. 10.1093/nar/gkaf763.

(61) Pezeshkian, W.; Ipsen, J. H. Creasing of Flexible Membranes at Vanishing Tension. Phys. Rev. E 2021, 103 (4), L041001. 10.1103/PhysRevE.103.L041001.

(62) Dasanna, A. K.; Fedosov, D. A. Mesoscopic Modeling of Membranes at Cellular Scale. Eur. Phys. J. Spec. Top. 2024, 233 (21), 3053–3071. 10.1140/epjs/s11734-024-01177-4.

(63) Duncan, A. L.; Pezeshkian, W. Mesoscale Simulations: An Indispensable Approach to Understand Biomembranes. Biophys. J. 2023, 122 (11), 1883–1889. 10.1016/j.bpj.2023.02.017.

(64) Siggel, M.; Kehl, S.; Reuter, K.; Köfinger, J.; Hummer, G. TriMem: A Parallelized Hybrid Monte Carlo Software for Efficient Simulations of Lipid Membranes. J. Chem. Phys. 2022, 157 (17), 174801. 10.1063/5.0101118.

(65) Harker-Kirschneck, L.; Hafner, A. E.; Yao, T.; Vanhille-Campos, C.; Jiang, X.; Pulschen, A.; Hurtig, F.; Hryniuk, D.; Culley, S.; Henriques, R.; Baum, B.; Šariæ, A. Physical Mechanisms of ESCRT-III–Driven Cell Division. Proc. Natl. Acad. Sci. 2022, 119 (1), e2107763119. 10.1073/pnas.2107763119.

(66) Sadeghi, M.; Noé, F. Large-Scale Simulation of Biomembranes Incorporating Realistic Kinetics into Coarse-Grained Models. Nat. Commun. 2020, 11 (1), 2951. 10.1038/s41467-020-16424-0.

(67) Davtyan, A.; Simunovic, M.; Voth, G. A. The Mesoscopic Membrane with Proteins (MesM-P) Model. J. Chem. Phys. 2017, 147 (4), 044101. 10.1063/1.4993514.

(68) Ayton, G. S.; Blood, P. D.; Voth, G. A. Membrane Remodeling from N-BAR Domain Interactions: Insights from Multi-Scale Simulation. Biophys. J. 2007, 92 (10), 3595–3602. 10.1529/biophysj.106.101709.

(69) Ma, Y.; Sun, S.; Huang, X.; Tian, L.; Li, L.; Wang, J. Membrane Vesicles Embedded with Multiple Curved Proteins Subjected to Osmotic Pressure. J. Mech. Phys. Solids 2025, 204, 106283. 10.1016/j.jmps.2025.106283.

(70) Ghazizadeh, E.; Zeidi, M.; Stroberg, W. Tubulation of Membrane Sheets by Curvature-Inducing Proteins. Biochim. Biophys. Acta BBA - Biomembr. 2026, 1868 (1), 184484. 10.1016/j.bbamem.2025.184484.

(71) Pezeshkian, W.; König, M.; Marrink, S. J.; Ipsen, J. H. A Multi-Scale Approach to Membrane Remodeling Processes. Front. Mol. Biosci. 2019, 6. 10.3389/fmolb.2019.00059.

