## Supplementary Figures and Notes for "Inward and Outward Tethers Read Out the Spontaneous Curvature of Cellular Membranes"

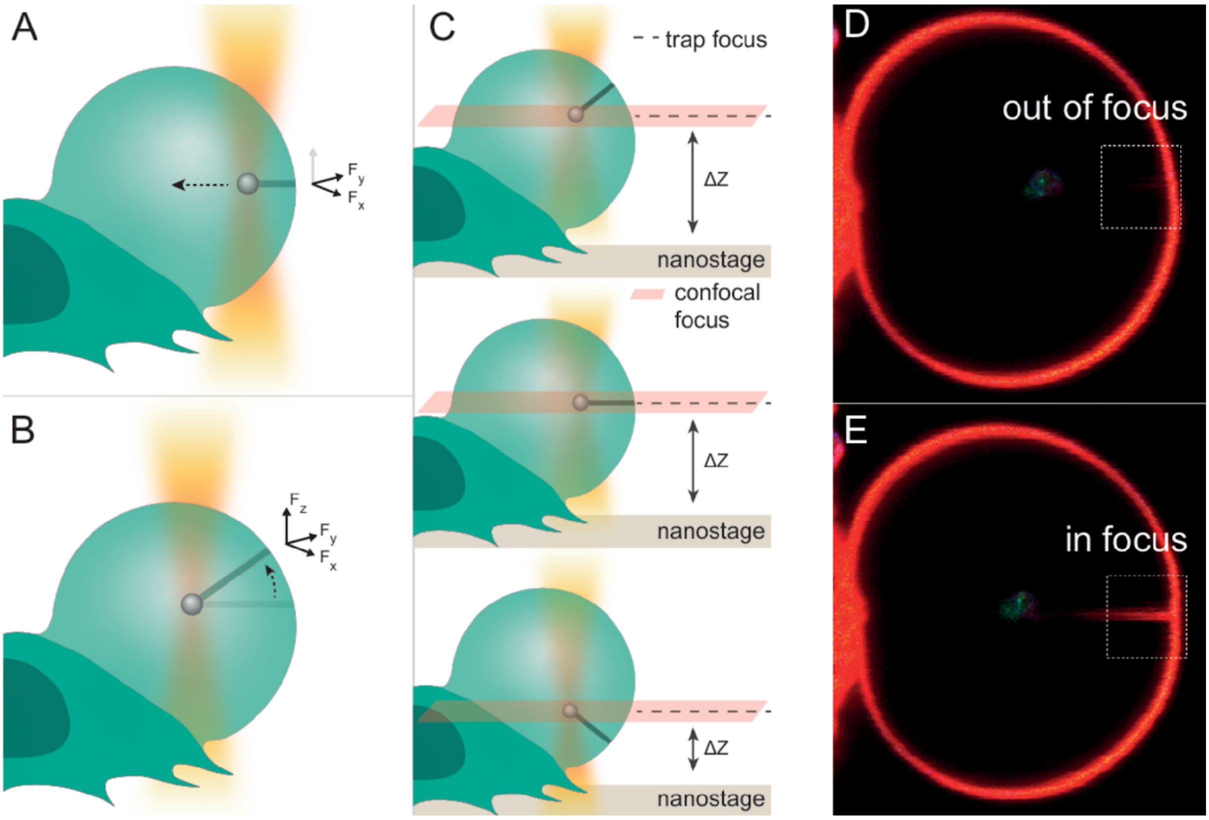

Supplementary Figure 1: **Z-component variability for in tether pulls.** (A) Schematic showing an in tether initially lying in the x–y plane. As the bead is pushed further into the GPMV, the tether slides upward along the spherical membrane (B), resulting in a new force direction with a z-component. (C) Confocal focus (red plane) and trap focus (dashed line) can be offset by adjusting the nanostage. This allows the tether to move above or below the focal plane while the trap remains in focus. (D, E) Confocal images of the same GPMV with an in tether appearing out of focus (D) and then brought into focus (E) by repositioning the nanostage and trap [adapted from: Effrosyni Drakouli. Quantifying Protein Crowding Effects driven by Mucins. MSc thesis, The Niels Bohr Institute, Faculty of Science, University of Copenhagen, 2025. Victoria Thusgaard Ruhoff. Reconstitution of viral budding. PhD thesis, The Niels Bohr Institute, Faculty of Science, University of Copenhagen, 2024.]

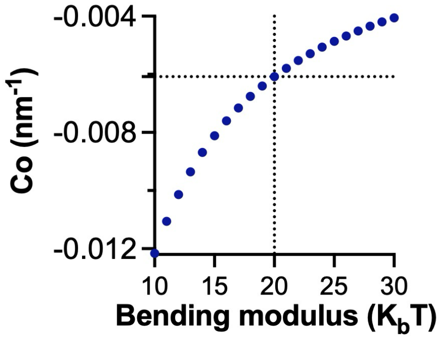

Supplementary Fig. 2: **Dependence of spontaneous curvature on bending rigidity at constant force difference $\Delta f$.** Here we use $\Delta f = 6.69\ pN$ from main text Fig. 1c and the literature bending modulus range (see main text section 1.1). The dashed lines show the value of $20\ k_BT$ often used throughout this work.

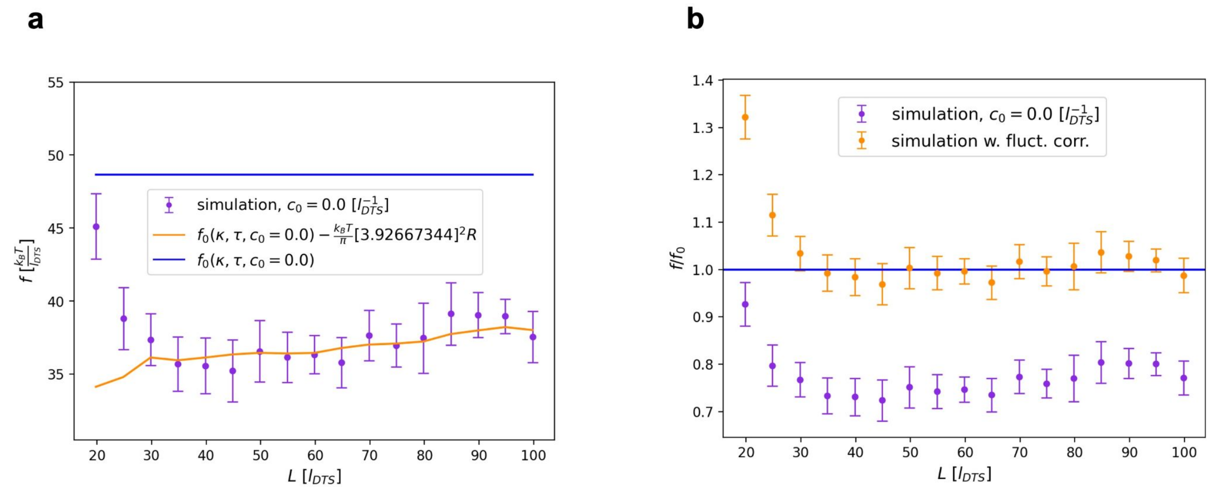

Supplementary Fig. 3: **Force Elongation Simulations.** The force pulling at different tether lengths is measured in simulations of a simple membrane. **(a)** Pulling force elongation curve and theoretical pulling force $f_0$without (blue) and with (orange) fluctuation correction $f_{0,corr} \approx f_0 - \frac{k_BT}{\pi}\Lambda^2 R$, the cut-off wavelength $\Lambda \approx 3.9\ l^{-1}{}_{DTS}$ was determined by a fit to the data (least squares). Corresponding to a cut-off length $2\pi/\Lambda \approx 1.6\ l_{DTS}$ this represents the resolution of FreeDTS simulations, as is very close to the average edge-link length ($1 l_{DTS} < edge < \sqrt{3}\ l_{DTS}$). **(b)** The ratio of measurements in simulations and theoretical prediction, without (purple) and with (yellow) the fluctuation correction applied to the theory solution. Each data point was obtained from configurations generated during the last $2 \times 10^6$MC steps of a long simulation of $10 \times 10^6$MC steps (see Methods section for more details).

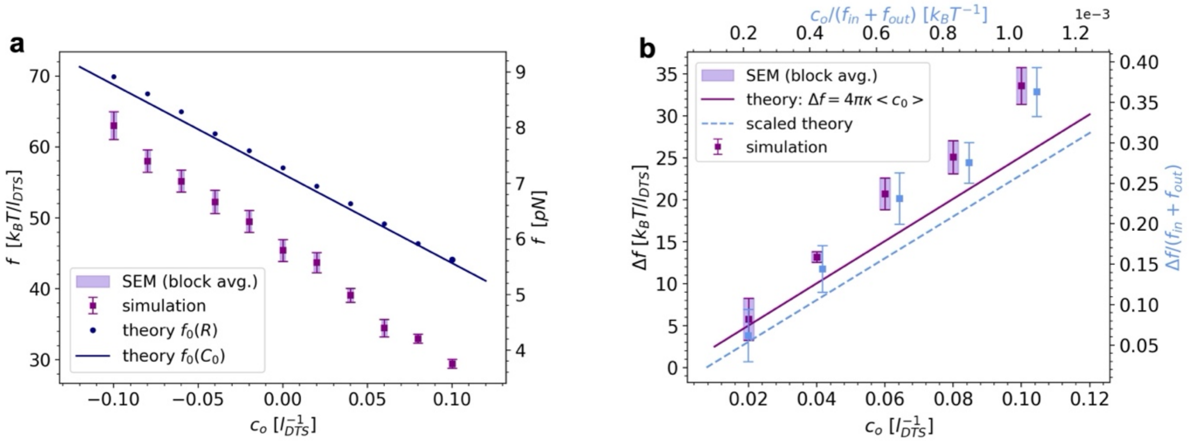

Supplementary Fig. 4: **Simulations of tether pulling with global homogeneous spontaneous curvature**. **(a)** Dependency of the pulling force on spontaneous curvature. **(b)** The difference between pulling forces measured on tethered membranes with the same positive vs. negative spontaneous curvature. Purple, left and bottom axis: no scaling; blue, right and top axis: scaled by the sum of in- and out-pulling force for comparability to experiments. All simulations were slowly pulled to the desired tether length (see Methods), then additional parameters such as spontaneous curvature were changed or added, and the simulation was restarted for another $10 \times 10^6$MC steps. The last $2 \times 10^6$MC steps were analysed.

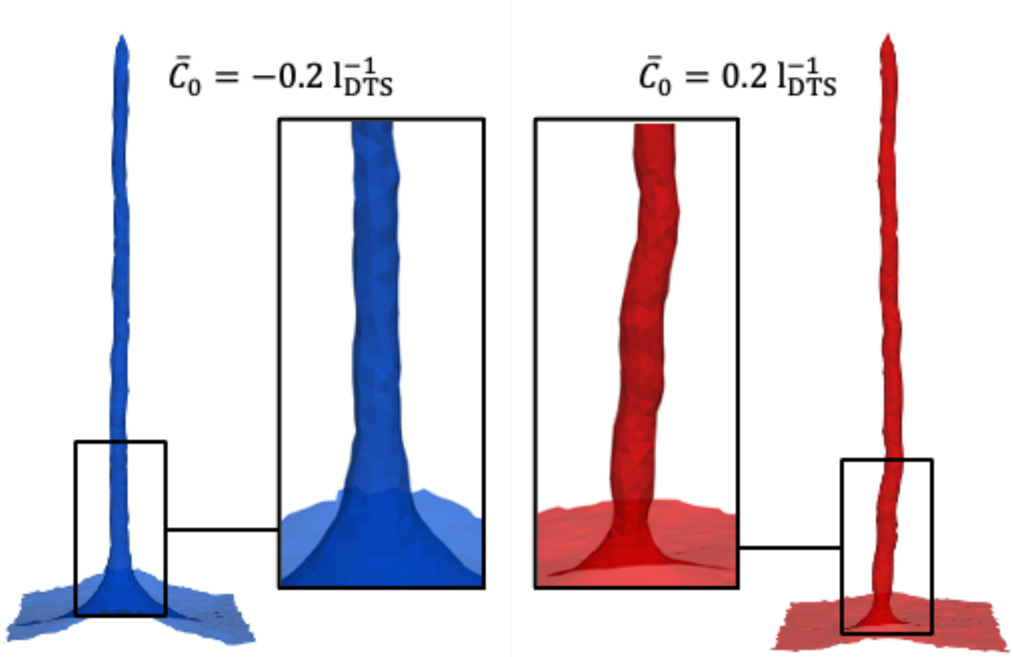

Supplementary Fig. 5: **Effect of spontaneous curvature $\bar{C}_0$ on membrane fluctuations on an out-tether.** At positive $\bar{C}_0$, which has the sign of the tether's innate curvature due to pulling out, membrane fluctuations are clearly visible, while at negative $\bar{C}_0$ (opposite sign of the tether's innate curvature) the fluctuations appear suppressed.

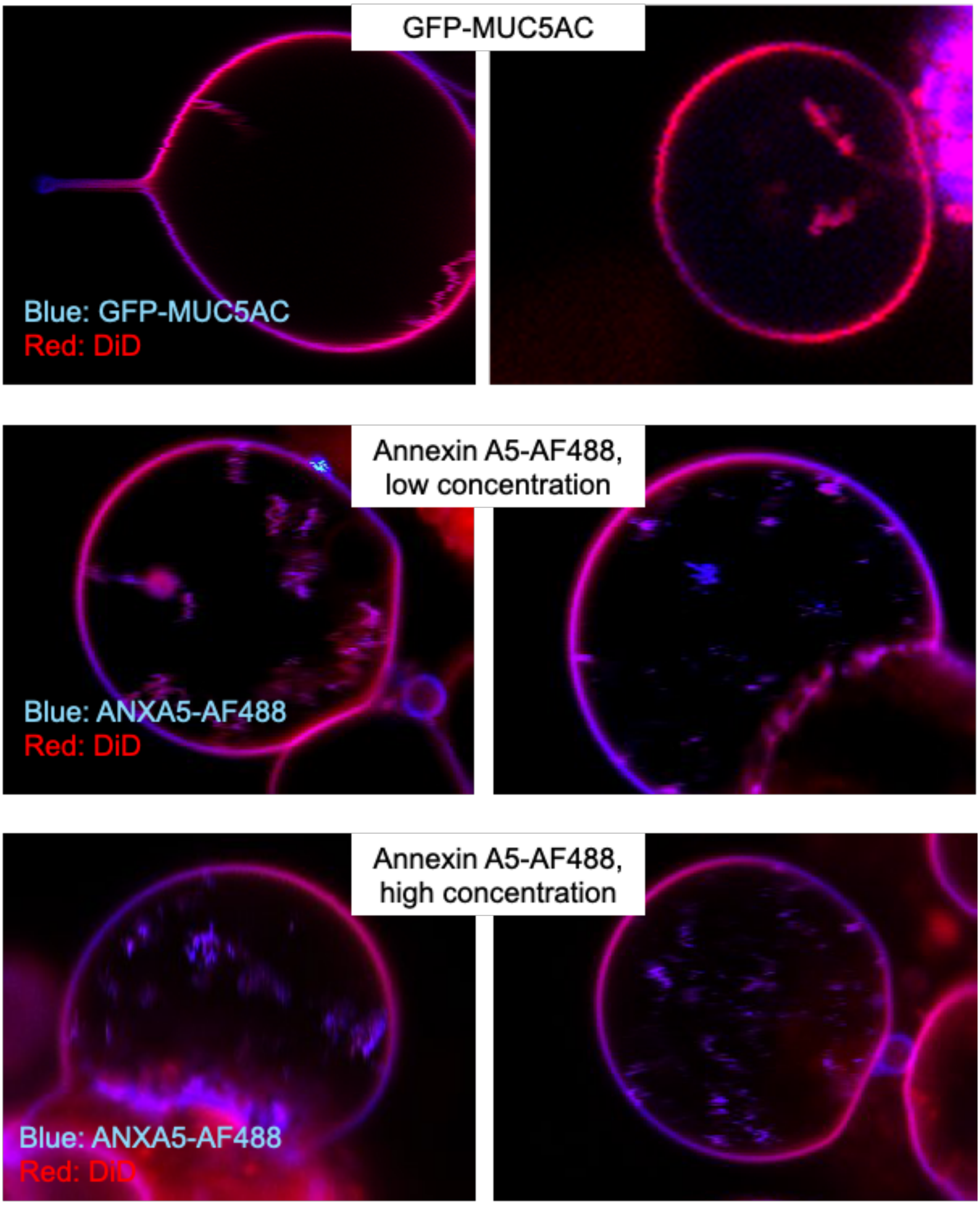

Supplementary Figure 6: **GPMVs from GFP-MUC5AC expressing cells (top) or extracellularly added Annexin A5-AF488 cells (bottom), where select examples displayed inner structures characteristic of negative curvature.**

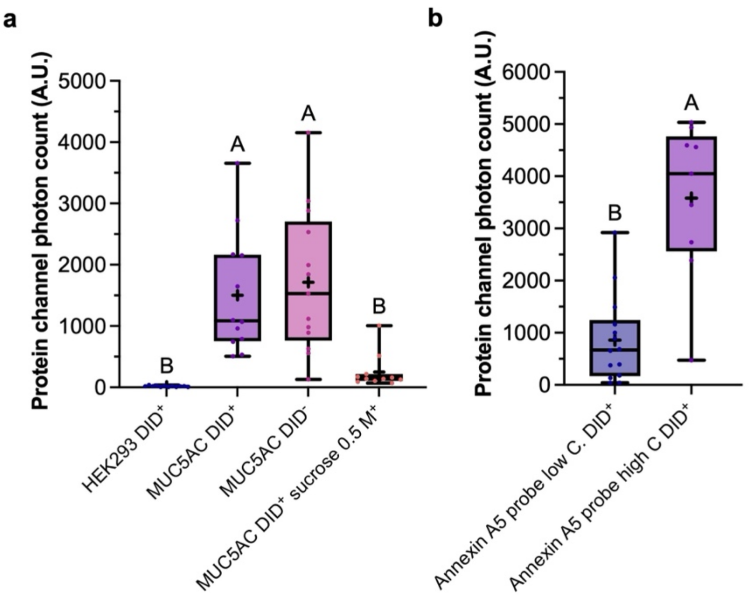

Supplementary Figure 7: **Protein density distributions for mucins and annexins.** Area below the fitted Gaussian distribution to the blue photon count, which is proportional to GFP-MUC5AC **(a)** or Annexin A5-AF488 **(b)** relative surface densities over the GPMV's analysed flat membrane regions. Number of experimental days, n, and datapoints, N: $n_{HEK}$ = 4, $N_{HEK}$ = 12; $n_{mucin}$ = 5, $N_{mucin}$ = 12; $n_{mucin\ no\ DiD}$ = 5, $N_{mucin\ no\ DiD}$ = 13; $n_{mucin\ sucrose}$ = 2, $N_{mucin\ sucrose}$ = 11; $n_{low\ C\ annexin}$ = 5, $N_{low\ C\ annexin}$ = 14; $n_{high\ C\ annexin}$ = 3, $N_{high\ C\ annexin}$ = 9.

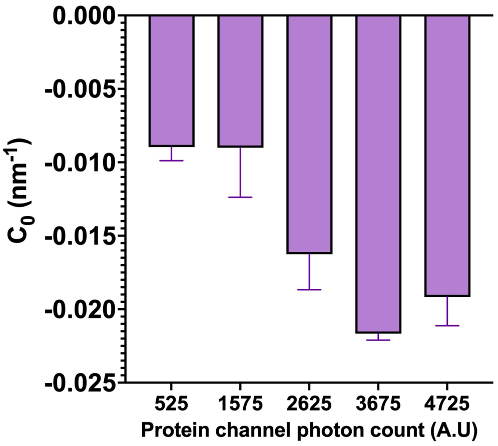

Supplementary Figure 8: **Addition of recombinant Annexin A5 to the external leaflet modulates the effective spontaneous curvature** ($nm^{-1}$) **in a concentration-dependent manner.** The protein-channel associated fluorescence at the GPMV surface is proportional to the protein surface density. $C_{0eff}$, calculated using equation 1 in the form $\Delta f = 4\pi\kappa C_{0eff}$, vs the area below the fitted Gaussian distribution to the blue photon count, which is proportional to the Annexin A5-488 relative surface densities over the GPMV "flat" membrane analyzed regions. Bin size: 1050 (A.U). Number of experimental days, n, and datapoints, N: $n_{low\ C\ annexin}$ = 5, $N_{low\ C\ annexin}$ = 14; $n_{high\ C\ annexin}$ = 3, $N_{high\ C\ annexin}$ = 9.

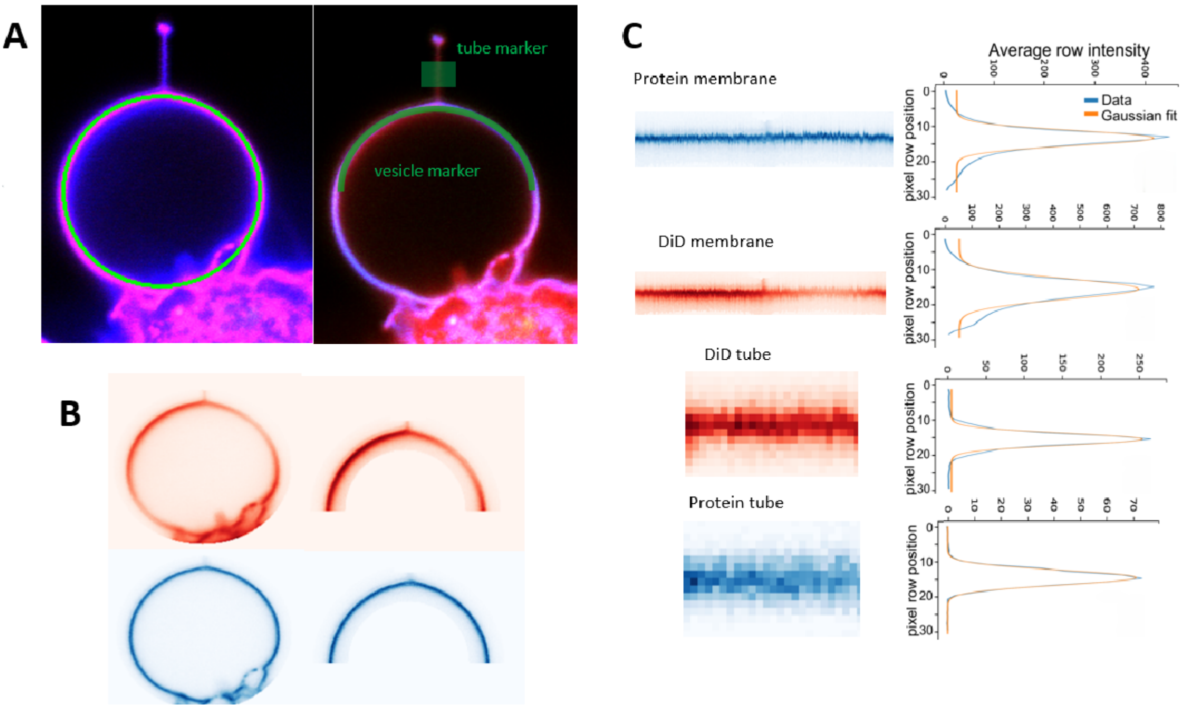

Supplementary Figure 9: **Data analysis for extracting relative tether size and curvature sorting of proteins.** (A) Confocal fluorescence images of a GPMV with an extended membrane tether, showing DiD (red) and protein marker (blue). The vesicle shape was manually approximated by fitting a circle (green), from which a 180° arc (upper hemisphere – vesicle marker) was selected with an arc width of 3.1 µm, centered at the fitted radius. A rectangular crop of width 3.1 µm (green rectangle-tube marker), centered along the line connecting two manually selected points at the tether ends, was also created for further analysis. (B) Top panels show the DiD channel, and bottom panels show the protein channel (GFP-MUC5AC, reference representative protein), for both the full vesicle (left) and the 180° cropped arc (right). (C) Intensity analysis of each region. For each rectangular crop (vesicle or tube-fixed size), pixel intensities were collapsed along columns (parallel to the membrane periphery or tether axis) by averaging values across rows. The resulting 1D intensity profiles are plotted as a function of pixel row position for both channels (blue: protein, red: DiD). These profiles were fitted with Gaussian functions (orange) to extract area below the curve, which are reported as the quantitative fluorescence intensity of the membrane-associated proteins and lipids [adapted from: Effrosyni Drakouli. Quantifying Protein Crowding Effects driven by Mucins. MSc thesis, The Niels Bohr Institute, Faculty of Science, University of Copenhagen, 2025. ]

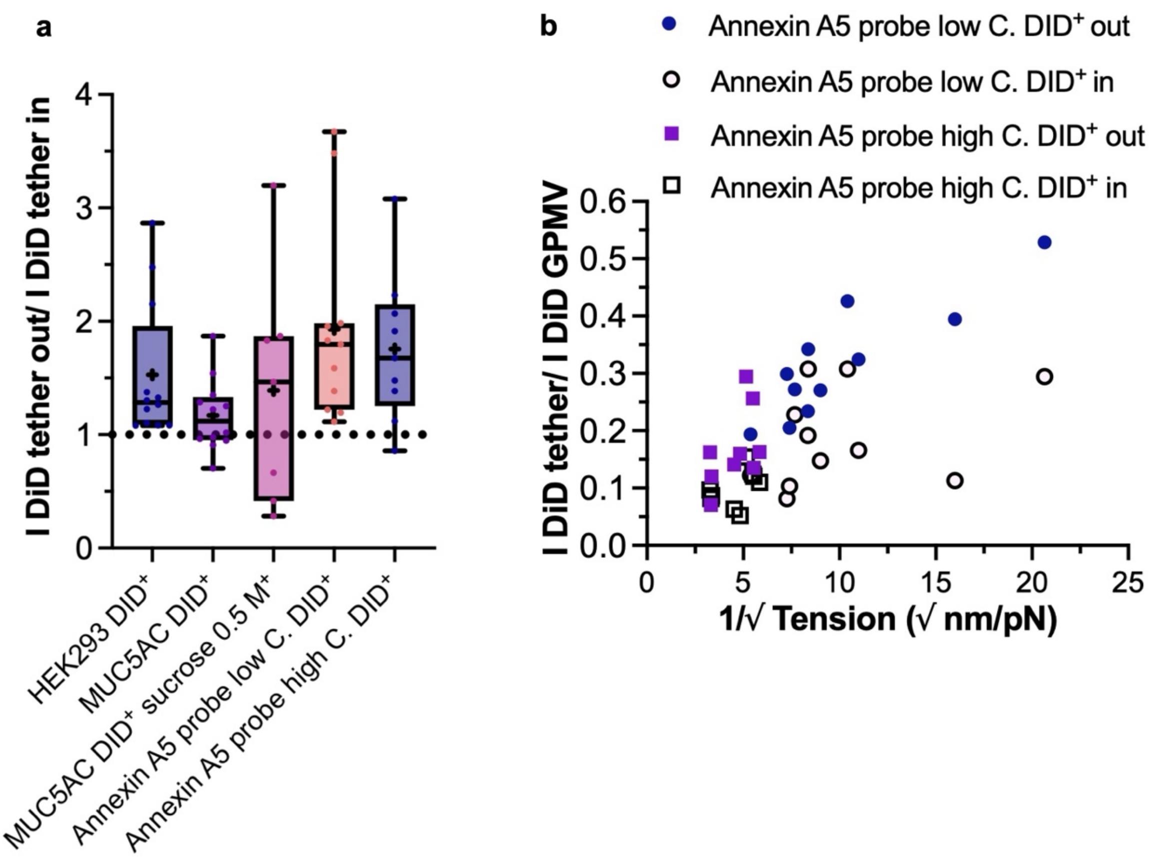

Supplementary Figure 10: **Relative tether radii. (a)** Intensity ratio of the membrane dye on in- vs. out-tethers from GPMVs derived from HEK293 cells under different conditions, showing them to be of similar size. This ratio is subject to out-of-focus and photobleaching effects, which are especially important for in-tethers, because they are hard to image in focus as explained in Supplementary figure 1, and we always do second the in tether, such that photobleaching effects increase. **(b)** Intensity ratio of the membrane dye on in-tethers or out-tethers with respect to the flat GPMV surface signal, which is proportional to the tether radius, vs. $\sqrt{1/\tau}$, corresponding to the tension shown in figure 2 D, from GPMVs derived from HEK293 cells under different conditions, showing the expected increase of "relative" tether radius (quantified from the confocal imaging) for an increasing $\sqrt{1/\tau}$**,** consistent with equation **S1-3**. A significant positive monotonic association was observed between $\sqrt{1/\tau}$ and the GPMV-tether intensity ratio under low annexin C out tethers (Spearman's $r_S$=0.80, exact two-sided P=0.0047). No statistically significant monotonic associations were detected under low annexin C in tethers ($r_S$=0.45, P=0.173), high annexin C out tethers ($r_S$=0.45, P=0.230), or high annexin C in tethers ($r_S$=0.52, P=0.162). Number of experimental days, n, and datapoints, N: $n_{HEK}$ = 4, $N_{HEK}$ = 12; $n_{mucin}$ = 5, $N_{mucin}$ = 12; $n_{mucin\ sucrose}$ = 2, $N_{mucin\ sucrose}$ = 7; $n_{low\ C\ annexin}$ = 5, $N_{low\ C\ annexin}$ = 11; $n_{high\ C\ annexin}$ = 3, $N_{high\ C\ annexin}$ = 9.

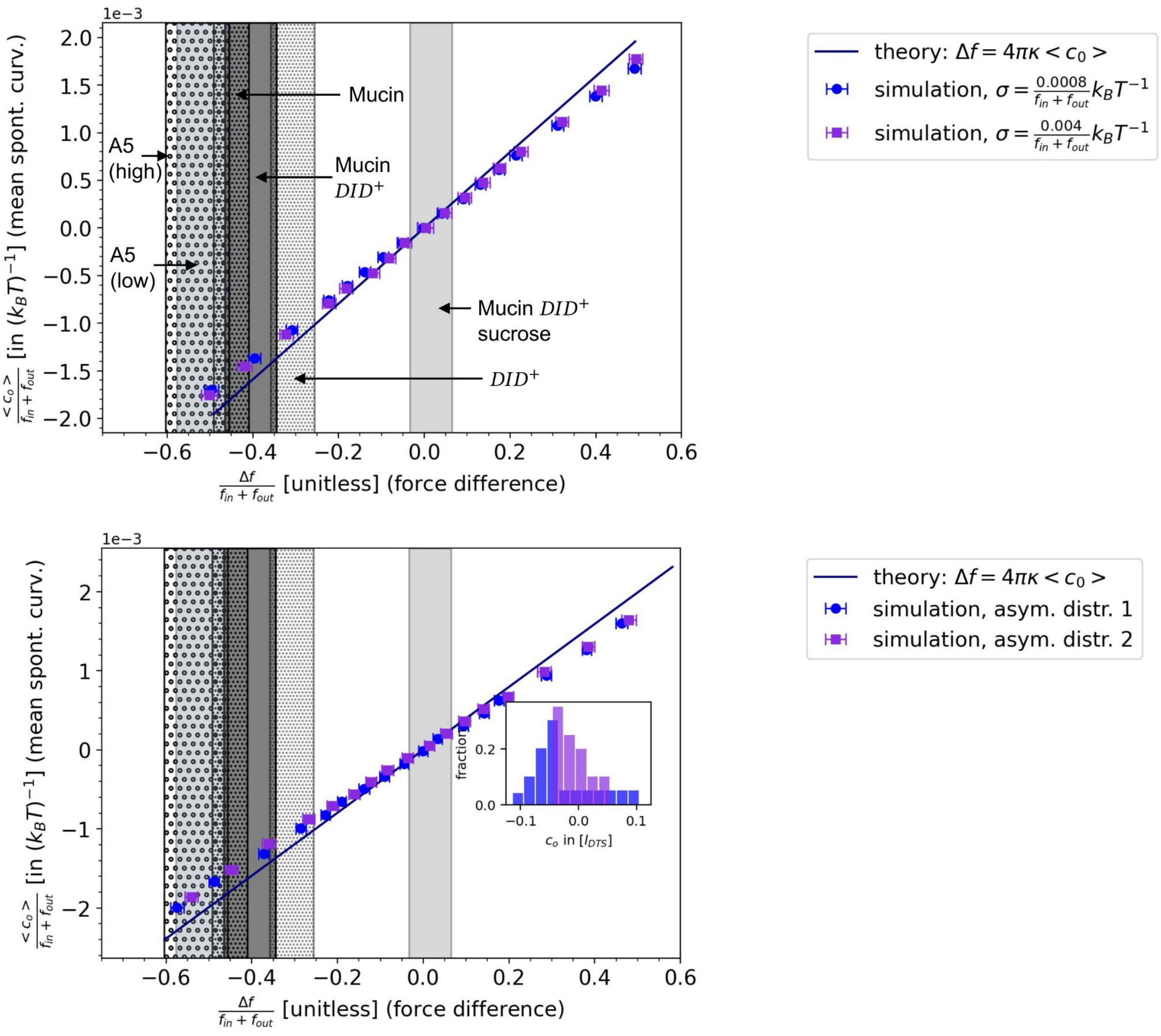

Supplementary Figure 11: **Force difference simulations of uniform (a) and asymmetric distributions (b).** Location of experimental measurements noted, as in main text Fig. 2f.

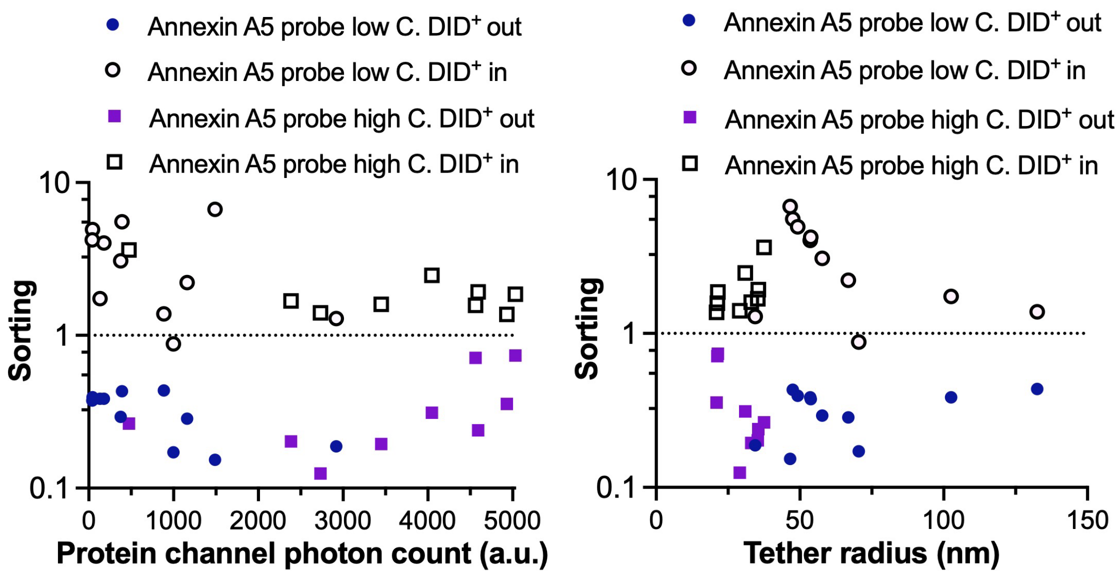

Supplementary Figure 12: **Experimental Annexin A5 sorting follows expected trends. (a)** Out/in Sorting vs. the area below the fitted Gaussian distribution to the blue photon count, which is proportional to the Annexin A5-488 relative surface densities over the GPMV "flat" membrane analyzed regions. The sorting increases with decreasing global GPMV protein density, as expected. **(b)** Out/in Sorting vs. the tether radius (nm), calculated using equation S1-3. For the low protein concentration, the in sorting increases with decreasing tether radius, as expected. However, the in sorting for the high protein concentration, and the out sorting for both concentrations, appear less dependent on the tether radius; consistent with an expected decreased sorting for higher global protein concentrations, and a depletion effect over the opposing preferred curvature. Number of experimental days, n, and datapoints, N: $n_{\text{low C annexin}} = 5$, $N_{\text{low C annexin}} = 11$; $n_{\text{high C annexin}} = 3$, $N_{\text{high C annexin}} = 9$.

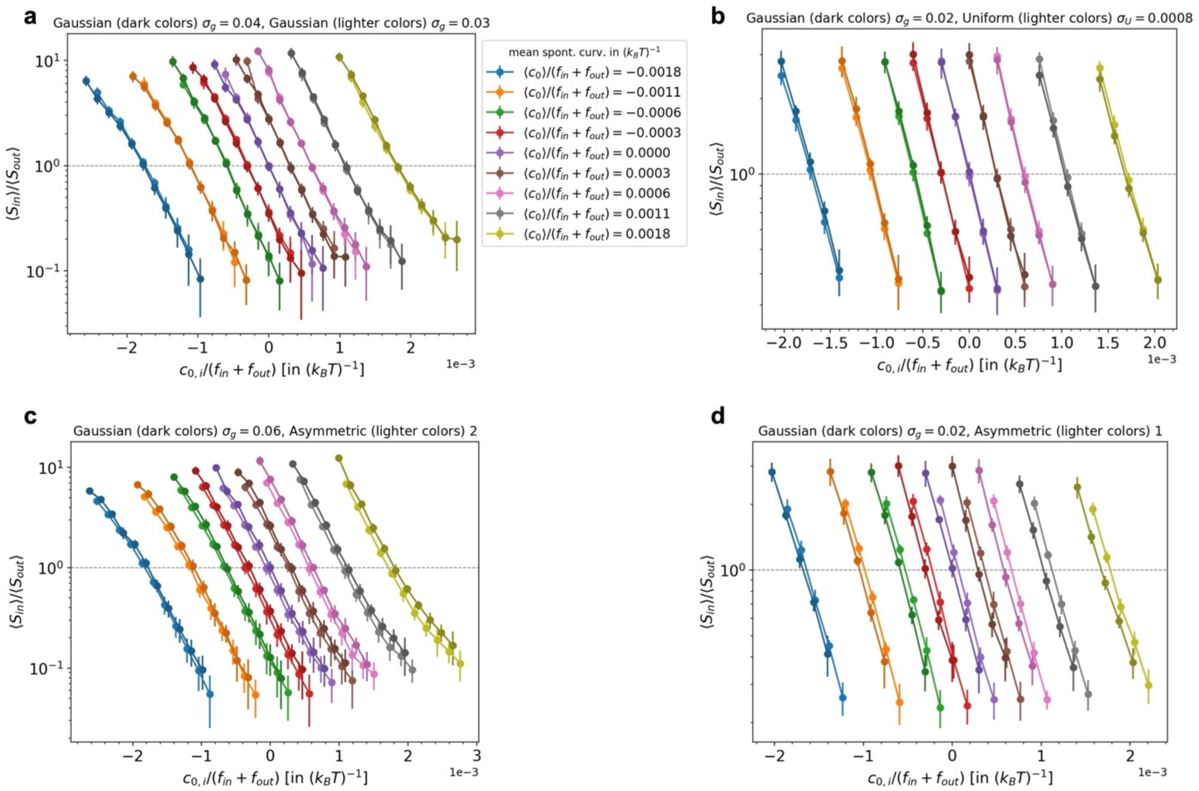

Supplementary Figure 13: **Inclusion sorting ratios from simulations for asymmetric distributions and distributions with narrower variances than those shown in the main text.**

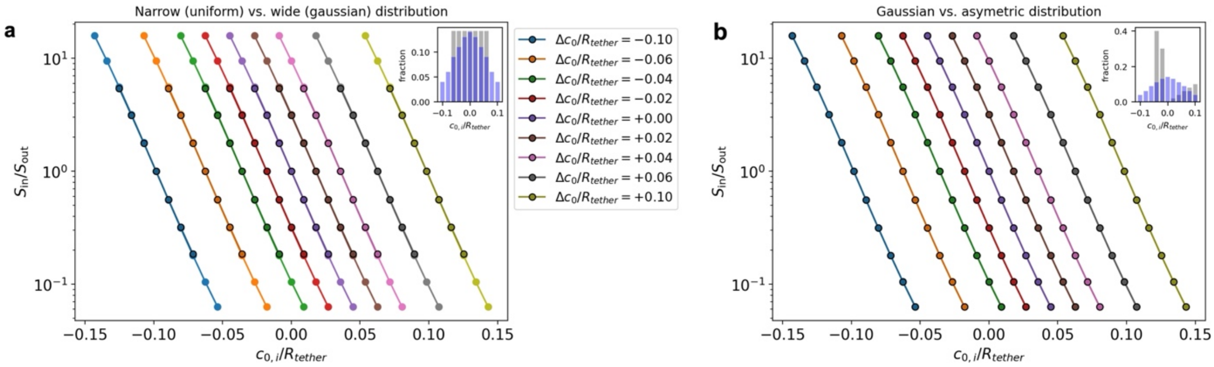

Supplementary Fig. 14: **Theoretical results on the sorting ratio on in- vs. out-tethers, derived from canonical ensembles for the flat and tethered membrane region with particle exchange between them at full coverage of the membrane with a heterogeneous curvature distribution. (a)** Results for different symmetric distributions, visualising that all symmetric distributions sort the same, independent of width, species number and type of distribution. **(b)** Results for a symmetric versus an asymmetric distribution, showing that while asymmetric distributions sort only approximately the same, for the curvature ranges in this work there is no distinguishable difference to symmetric distributions.

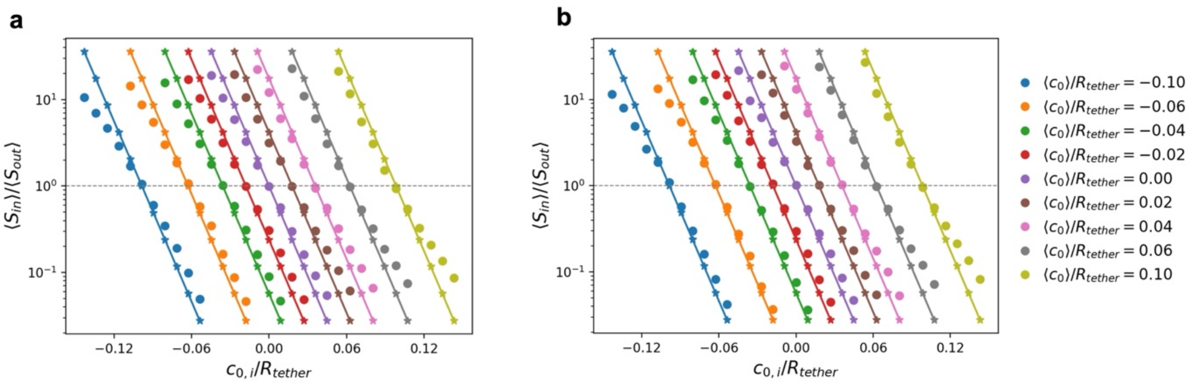

Supplementary Fig. 15: **Simulated sorting (points) versus theoretical results (lines) with input of the theoretical radius $R_{in/out} = R_{tether} = \sqrt{\kappa/2\tau}$ (Eq. 2, main text). (a)** Results for the widest uniform distribution. **(b)** Results for the widest gaussian distribution. In both plots, the curvature axis is scaled by the theoretically expected tether radius $R_{tether}$.

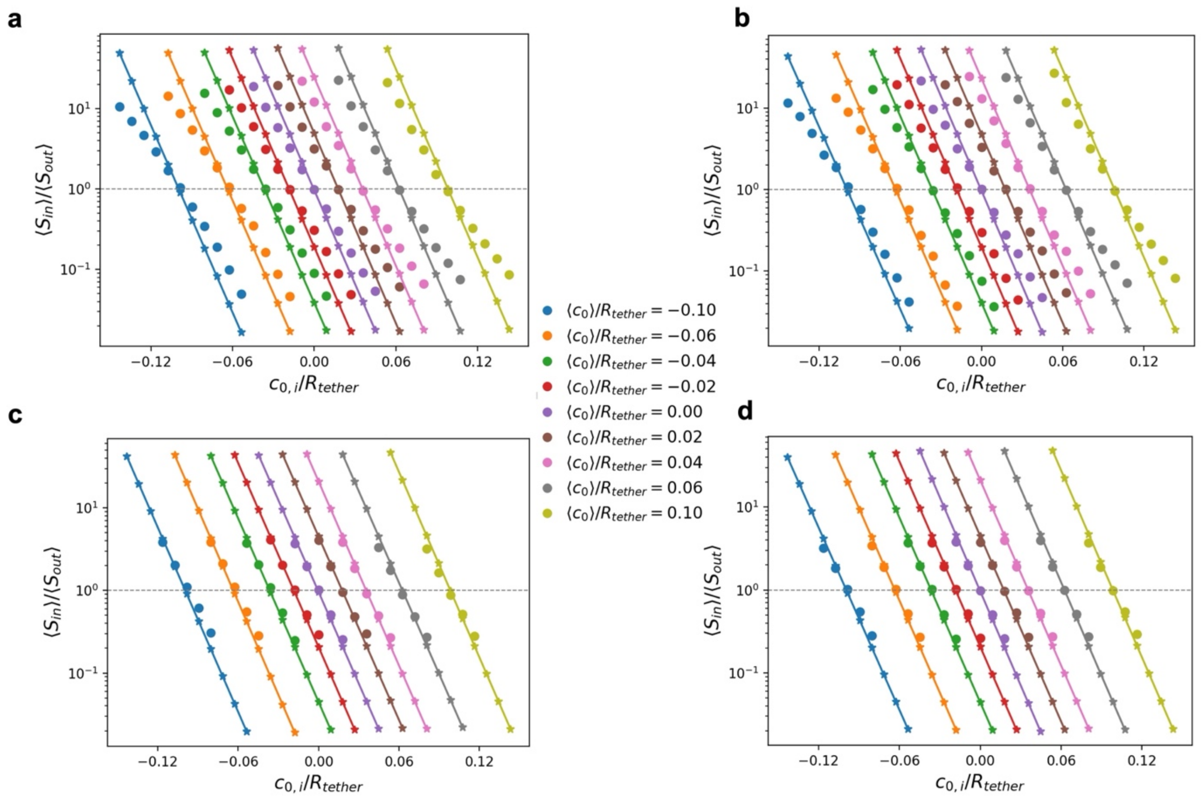

Supplementary Fig. 16: **Simulated sorting (points) versus theoretical results (lines) with input of the radii $R_{sim,in}$, $R_{sim,out}$ from simulations. (a&c)** results for the widest (a) and narrowest (c) uniform distributions with $\sigma = 4/R_{tether} \times 10^{-3}$and $\sigma = 8/R_{tether} \times 10^{-4}$ respectively. **(b&d)** results for the widest (b) and narrowest (d) gaussian distributions with $\sigma = 0.06/R_{tether}$and $\sigma = 0.02/R_{tether}$ respectively. In both plots, the curvature axis is scaled by the theoretically expected tether radius $R_{tether} = \sqrt{\kappa/2\tau}$ (Eq. 2, main text).

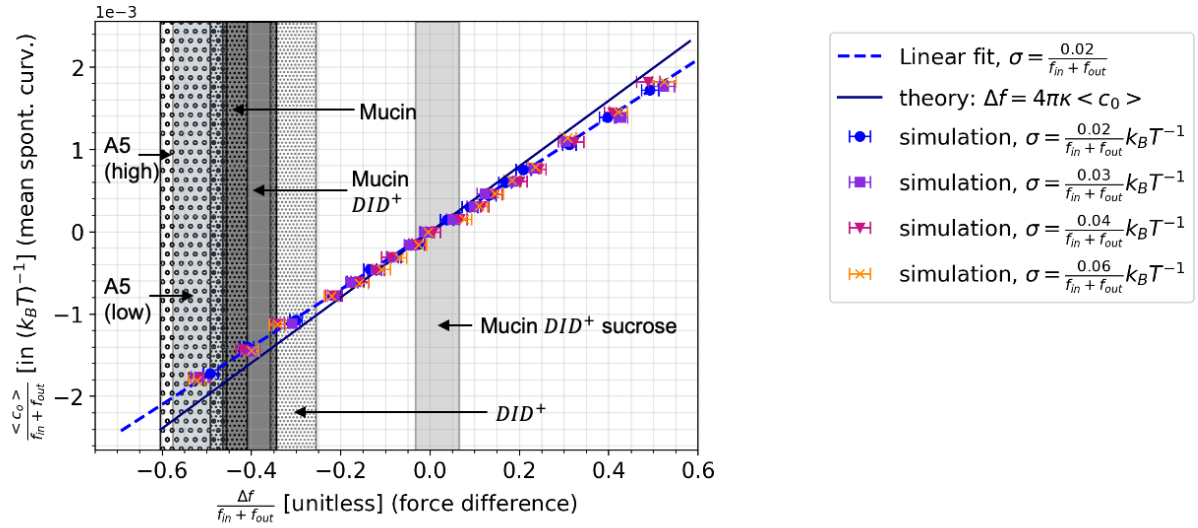

Supplementary Fig. 17: **Read off version of Fig. 2f (main text) of spontaneous curvature as a function of force difference based on simulations of gaussian inclusion distributions.**

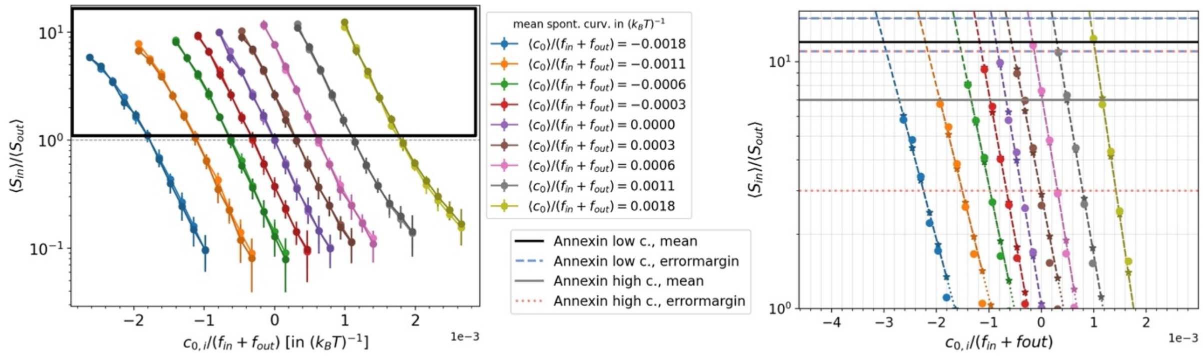

Supplementary Fig. 18: **Read off version of Fig. 3c (main text) of curvature-induced sorting as a function of spontaneous curvature based on simulations of gaussian inclusion distributions.**

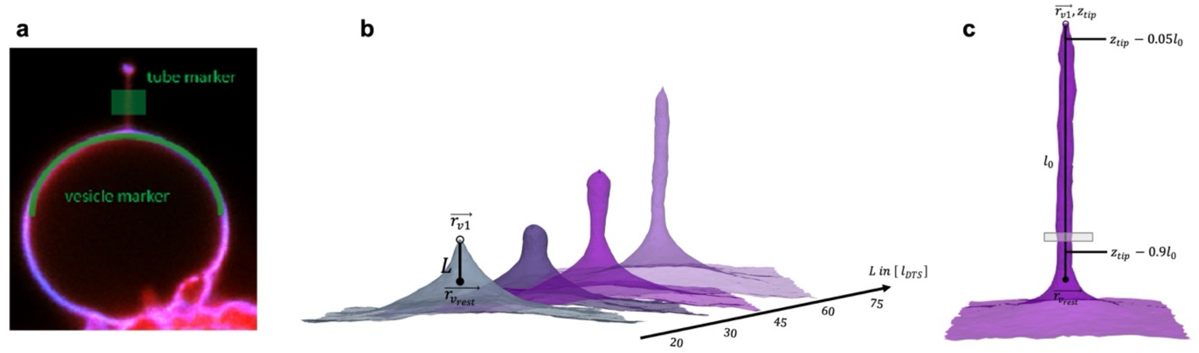

Supplementary Fig. 19: **Visualization of Methods: Analysis of protein sorting in experiments (a) and process of tether pulling & radius analysis in simulations (b&c)**. (a) Confocal fluorescence images of a GPMV with an extended membrane tether, and an overlay of DiD membrane marker (red) and protein marker (blue). **(b)** Tethers are pulled from a flat membrane in PBC via a harmonic potential between a chosen tip vertex (position $\overrightarrow{r_{v1}}$ ) close to the center of the box and the other vertices (center of mass at $\overrightarrow{r_{v_{rest}}}$). The desired distance $L$ minimizing the potential is slowly increased to pull tethers of different lengths. These are subsequently equilibrated further (for $8 - 12 \times 10^6$ MC steps) at their $L$ before analysis in simulations with additional parameters and/or proteins. **(c)** Analysis of tether radius. Procedure of determining the tether radius as the mean over the radius recovered from slices (gray box) along the z-axis within the limits of 5-90% of the tether length $l_0$below the tether's tip vertex.

### Supplementary Note 1

#### Analytical Model for Membrane Tether Pulling

The bending energy of a tether pulled from a flat membrane patch can be described by the Helfrich Hamiltonian,

$$E_B = \oint \left[\frac{\kappa}{2}(2H - \bar{C}_0)^2 - \kappa_G K\right] dA \qquad \text{(S1-1)},$$

with bending modulus $\kappa$, gaussian modulus $\kappa_G$ and spontaneous curvature $\underline{C}_0$. The Gaussian term (second term) is constant if there is no topological change to the surface and therefore will be neglected. For an idealized thin membrane tether of radius $R$ and length $L$, contributions from the small tether tip and base regions become negligible and the bending energy including spontaneous curvatures follows as the sum of the integral over the semi-flat part and the tether region:

$$E_B = \oint \frac{\kappa}{2}\bar{C_0}^2 dA_{flat} + \int_0^L \pi R\kappa\left(\frac{1}{R} - \bar{C}_0\right)^2 ds = \frac{\kappa}{2}\bar{C_0}^2(A_{tot} - 2\pi RL) + \pi L\kappa\left(\frac{1}{R} - 2\bar{C}_0 + \bar{C}_0^2 R\right) \qquad \text{(S1-2)}.$$

The total energy of the system furthermore must account for the pulling of the tether as well as the membrane tension. We also include a term constraining the surface area $A_{tot}$ of the membrane to a constant $A_0$ via an area compressibility $K_A$, i.e $E_A = K_A(A_{tot} - A_0)^2$. Therefore $E_{tot} = E_B + E_A + E_\tau + E_{pull}$ with the pulling force $f$ leading to a contribution $E_{pull} = fL$ and an external tension applied on a membrane patch leading to $E_\tau = -\tau A_p = -\tau(A_{tot} - 2\pi RL)$. Please note: this external frame tension ensemble treats tension as the mechanical work performed on a system but is equivalent to the often used treatment of tension $\sigma$ as the internal stresses from membrane area stretching [28]. Our frame tension ensemble reproduces the expected relations between internal and external stresses (for details see [64]). The radius of a stable tether will then be:

$$0 = \frac{\partial E_{tot}}{\partial R} \Rightarrow R = \sqrt{\frac{\kappa}{2\tau}} = R|_{\bar{C}_0 = 0} \qquad \text{(S1-3)}.$$

The minimal required force for tether extrusion, which is then the holding force, must satisfy $f \geq \frac{\partial(E_B + E_\tau)}{\partial L}$. Hence,

$$f_0 = \pi\kappa\left(\frac{1}{R} - 2\bar{C}_0\right) + 2\pi\tau R = 2\pi\left(\sqrt{2\tau\kappa} - \kappa\bar{C}_0\right) \qquad \text{(S1-4)}.$$

A simple, but powerful relationship then arises between the holding force for a membrane with spontaneous curvature $-\bar{C}_0$ and another one with $\bar{C}_0$:

$$\Delta f = f_0(-\bar{C}_0) - f_0(\bar{C}_0) = f_{in} - f_{out} = 4\pi\kappa\bar{C}_0 \qquad \text{(S1-5).}$$

$$f_{in} + f_{out} = 4\pi\sqrt{2\tau\kappa} \Leftrightarrow \tau = \frac{(f_{in}+f_{out})^2}{32\,\pi^2\kappa} \qquad \text{(S1-6).}$$

This relation (S1-5) theoretically provides a tool to measure membrane curvature experimentally via tether pulling experiments. However, experiments often underlie additional complexity, be it membrane fluctuations, for which a correction to Eq. (S1-4) is available [11], or the fact that multiple proteins and lipid types may be present, especially in realistic membranes, making it unclear what $\bar{C}_0$ represents exactly. In this work we demonstrate that Eq. (S1-5) is also a robust tool that can be reliably applied to several different, complex scenarios of membrane curvature.

While this widely used theoretical model assumes that the top and bottom of the tether are negligible, one can also include them explicitly in the calculations. With the top modelled as a spherical cap (half of a sphere) and the bottom neck that fuses the tether to the flat membrane modelled as the lower, inner quarter of a torus:

$$E_B = \oint \frac{\kappa}{2}\bar{C}_0^{\,2} dA_{flat} + \int_0^L \pi R\kappa\left(\frac{1}{R} - \bar{C}_0\right)^2 ds + \int_0^\pi d\phi \int_0^\pi \frac{\kappa}{2}\left(\frac{1}{R} - \bar{C}_0\right)^2 R^2 sin(\theta) d\theta +$$

$$\int_0^{2\pi} d\theta \int_\pi^{3\pi/2} \frac{\kappa}{2}\left(\frac{2}{R}\left(\frac{1+cos(\phi)}{2+cos(\phi)}\right) - \bar{C}_0\right)^2 R^2\left(2 + cos(\phi)\right) d\phi$$

$$= \frac{\kappa}{2}\bar{C}_0^{\,2}(A_{tot} - 2\pi RL) + \pi L\kappa\left(\frac{1}{R} - 2\bar{C}_0 + \bar{C}_0^2 R\right) + \pi\kappa(1 - \bar{C}_0 R)^2$$

$$+\pi\kappa\bar{C}_0^2 R^2(\pi - 1) - 2\pi\kappa\bar{C}_0 R(\pi - 2) \qquad \text{(S1-7).}$$

In this case, while the force difference $\Delta f$ will remain unaffected by the new contributions that account for the limited system size, the holding force itself as well as the radius of the tether will change. The minimal force required to pull a tether remains unaffected in its dependence on the radius, since terms including the cap and bottom of the tether do not depend on the length $L$. So still:

$$f_{min} = \pi\kappa\left(\frac{1}{R} - 2\bar{C}_0\right) + 2\pi\tau R \qquad \text{(S1-8).}$$

However, the equilibrium radius does change. It can then be determined from

$$0 = \frac{\partial E_B}{\partial R} = \frac{1}{R^2} - R\frac{2\bar{C}_0^2}{L}(\pi - 1) + 2\frac{\bar{C}_0}{L}(\pi - 1) - 2\frac{\tau}{\kappa} \qquad \text{(S1-9)}$$

and it will now clearly depend on both the spontaneous curvature as well as the length of the tether. This equation requires numerical treatment for solutions. Supplementary Figure S20 shows the right-hand side of Eq. (9) in the range of experimentally measured tensions and spontaneous curvatures. Then the equilibrium radius and force are recovered as functions of the spontaneous curvature and depending on tether lengths in Supplementary Figure S21.

It is hence important to conduct simulations and experiments in a regime, where the cap and bottom of the tether are negligibly small compared to the tether length, so that there is no significant dependence of the equilibrium radius and pulling force on the spontaneous curvature and tether length. Such a condition is roughly fulfilled when $\frac{R}{L} \leq 0.03$ as can be estimated from Supplementary Figure S21.

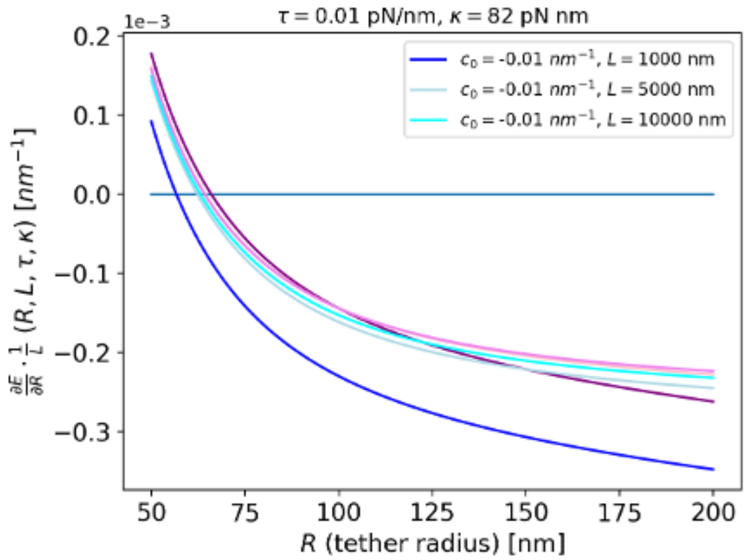

Supplementary Fig. 20: Derivative of the bending energy as a function of tether radius for a tether with a spherical cap tip and toroidal neck bottom to determine its equilibrium radius for different spontaneous curvatures of the membrane (equation (S1-9)). The intersection of the light blue horizontal line with the curves marks the solution of equation S1-8, i.e. the radii that minimize the bending energy.

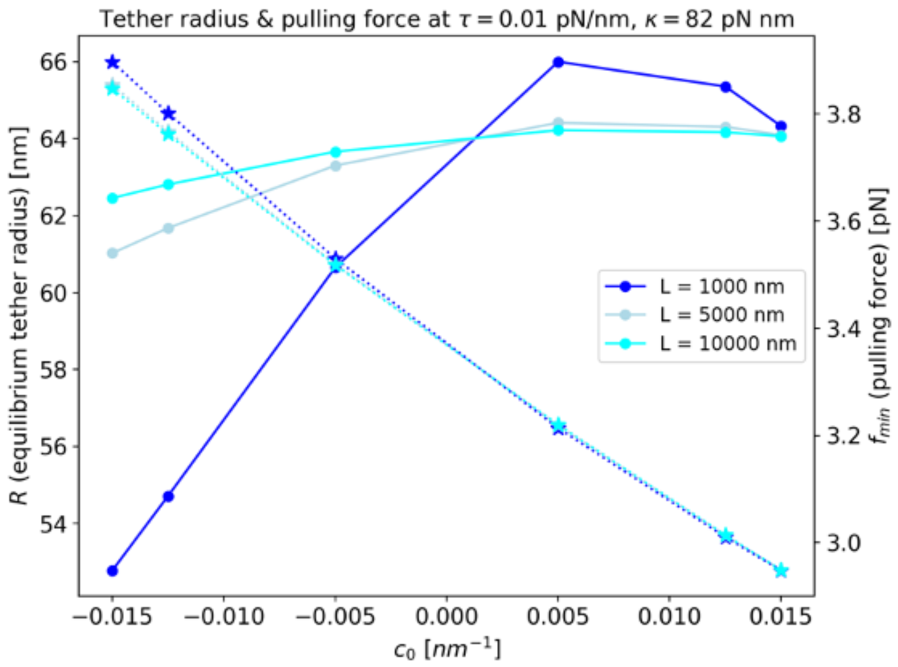

Supplementary Fig. 21: Equilibrium radii and holding forces, i.e. minimal pulling forces for tethers with spontaneous curvature described by cylinders with a spherical cap tip and toroidal neck bottom, i.e. solutions of eq. (S1-9).

### Supplementary Note 2

#### Simulated dual tethers with non-zero spontaneous curvature

Using tethers with $L = 60l_{DTS}$ safely within the force plateau, we tested the effect of global, homogeneous spontaneous curvature in our model (Eq. 3 with $\bar{C}_0 \neq 0$). The force measured in simulations had a similar qualitative dependence on spontaneous curvature as the theoretical prediction (Eq. 3) in the tested range $-0.1{l_{DTS}}^{-1} \leq \bar{C}_0 \leq 0.1{l_{DTS}}^{-1}$ ( $0.1{l_{DTS}}^{-1} \simeq 5 \times 10^{-3}\ nm^{-1}$, see Methods). Similar to the $\bar{C}_0 = 0$ case forces remained 10%-30% below the theory without fluctuation correction, deviating less for negative $\bar{C}_0$ and more for positive $\bar{C}_0$ on an out-tether with inherent positive curvature (Supplementary Figure S4a). Interestingly, the force difference $\Delta f$ (Eq. 1) showed closer agreement to the theoretical expectation (Supplementary Figure S4b). Especially at weaker curvatures $|\bar{C}_0| \leq 0.05\sqrt{2\tau/\kappa}$ it was accurate without a fluctuation correction, suggesting that $\Delta f$ inherently cancels deviating factors and is more robust than the holding force $f$. Yet, $\Delta f$ was still affected by the different influence of positive vs. negative $\bar{C}_0$ on membrane fluctuations. The required amount of fluctuation correction on $f$ varied with $\bar{C}_0$ (on out-tethers: free fluctuations at $\bar{C}_0 > 0$, strong fluctuation correction needed vs. squeezing effect suppressing fluctuations at $\bar{C}_0 < 0$, less fluctuation correction needed; see Supplementary Figure S5). At larger absolute spontaneous curvatures this deformation of tether from its idealized smooth cylinder shape also caused $\Delta f$ to deviate from Eq. 1.

To summarize, the mesoscale simulations
**a)** reproduce the theoretical tether pulling force for simple membranes at $\bar{C}_0 = 0$,
**b)** show that at $\bar{C}_0 \neq 0$, $\Delta f$ agrees better with theory than $f$ by partially compensating the effect of membrane fluctuations,
**c)** allow direct comparison to experiments via rescaling to $\Delta f/(f_{in} + f_{out})$.

### Supplementary Note 3

#### Theoretical description of curvature sorting

In this note, we derive an analytical expression for the sorting index (the ratio of sorting to inward vs. to outward membrane tethers) using a simple lattice-gas model for membrane inclusions with curvature-dependent interactions.

Let us divide the membrane surface into area elements of equal size $a_0$, and assign one inclusion to each element. Each inclusion type $i$ is characterized by a spontaneous curvature $c_{0,i}$. The membrane surface available to the inclusions is divided into two compartments:
(1) a tube with constant radius, $R$ and total area of $A_1$, and total inclusion number $M_1 = A_1/a_0$.
(2) a flat membrane reservoir with total area of $A_2$, and total inclusion number $M_2 = A_2/a_0$.
Therefore, the total number of inclusions will be $M = M_1 + M_2$.

The system consists of $I$ inclusion types. For each type $i$, $N_i$ inclusions are present, with $N_{i1}$ inclusions located in the tubular compartment and $N_{i2}$ in the flat compartment, such that$N_i = N_{i1} + N_{i2}$.

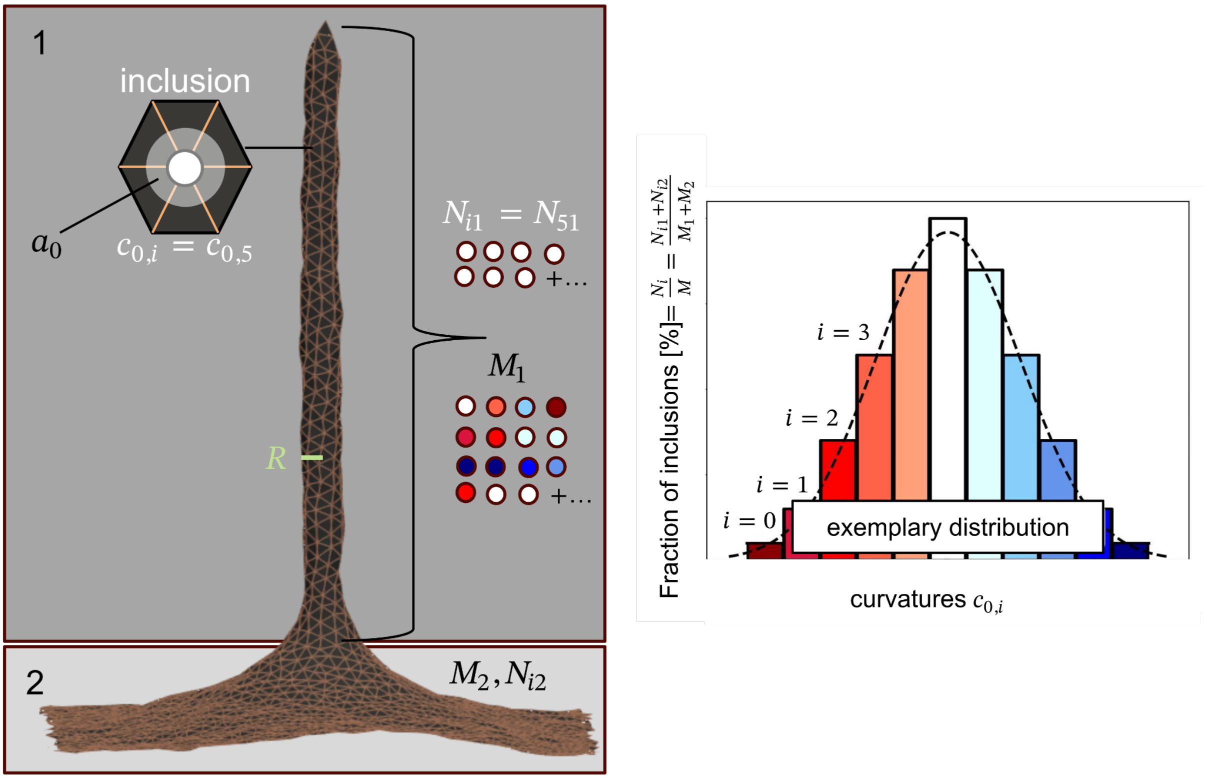

Supplementary Fig. 22: Schematic of theory model of curvature-induced inclusion sorting.

**Energy of the system**

Since the spontaneous curvature of the inclusion $i$ is $c_{0,i}$, the bending energy of a single inclusion of type $i$ in compartment 1 is $e_{i1} = -2\kappa c_{0,i}\frac{a_0}{R}$ and $e_{i2} = 0$ in the second compartment (Note that these expressions are obtained from the Helfrich Hamiltonian after choosing the flat membrane as the zero-energy reference state). Therefore, the total energy of the system system becomes $E = \sum_{i=0}^{I} e_{i1} N_{i1}$.

**Mixing Entropy**

The mixing entropy will be: $S = k_B ln(\Omega_1 \Omega_2)$ with the numbers of microstates

$$\Omega_1 = M_1!/\prod_{i=0}^{I} N_{i1}! \text{ and } \Omega_2 = M_2!/\prod_{i=0}^{I}(N_i - N_{i1})! \quad \text{(S3-1)}.$$

Using the Stirling approximation, the mixing entropy is obtained as:

$$S = k_B[M_1 ln(M_1) + M_2 ln(M_2)] - k_B[\sum_{i=0}^{I} N_{i1} ln(N_{i1}) + \sum_{i=0}^{I}(N_i - N_{i1}) ln(N_i - N_{i1})] \quad \text{(S3-2)}.$$

**Free Energy**

The system free energy is defined as $F = E - TS$, and therefore:

$$F \approx \sum_{i=0}^{I} e_{i1} N_{i1} - \beta^{-1}[M_1 ln(M_1) - \sum_{i=0}^{I} N_{i1} ln(N_{i1}) + M_2 ln(M_2) - \sum_{i=0}^{I}(N_i - N_{i1}) ln(N_i - N_{i1})] + const. \quad \text{(S3-3)},$$

where $\beta = 1/k_B T$. We impose the constraint that the total number of the inclusions is also constant, since the membrane should be fully covered and the number of membrane elements should be constant, i.e.

$$M_1 = \sum_{i=0}^{I} N_{i1} \quad \text{(S3-4)}.$$

We can then minimize our free energy under this constraint. Using a Lagrange multiplier $\phi$ we minimise the free energy with respect to the free energy variables ($N_{i1}$s) for all species $i$.

$$\underline{F} = F + \phi(M_1 - \sum_{i=0}^{I} N_{i1}) \quad \text{(S3-5)}$$

$$\frac{\partial \underline{F}}{\partial N_{i1}} = e_{i1} + \beta^{-1}[ln(N_{i1}) - ln(N_i - N_{i1})] - \phi = 0 \quad \text{(S3-6)}$$

This leads to

$$N_{i1} = N_i/[1 + exp(\beta(e_{i1} - \phi))] \quad \text{(S3-7)}$$

Plugging eq. (S3-7) into eq. (S3-4) yields the link between the Lagrange multiplier $\phi$ to constant parameters of the system:

$$M_1 = \sum_{i=0}^{I} N_{i1} = \sum_{i=0}^{I} N_i/[1 + exp(\beta(e_{i1} - \phi))] \quad \text{(S3-8)}$$

This shows that $\phi$ depends on $M_1$ and $N_i$'s and can be solved numerically knowing the membrane tether size, and the underlying distribution of the inclusion type population. It represents competition between the species due to curvature preference and limited space.

In experiments, the flat membrane region 2 (GPMV) is significantly larger than the tether region 1. Hence, the density of a protein species on the flat region will not measurably decrease even if that species sorts strongly to the tether. Hence, we need to describe sorting relative to the global, constant density of a species, i.e. $S_{in/out,i} = \frac{\rho_{tether}}{\rho_{global}} = \frac{N_{i1}/M_1}{N_i/(M_1+M_2)}$ (S3-9).

As the sorting of a species to the out-tether one therefore finds:

$$S_{out,i} = \frac{N_{i1}/M_1}{\rho_{i,global}} = \frac{M_1+M_2}{M_1}\frac{N_{i1}}{N_i} = \frac{M_1+M_2}{M_1}\frac{1}{1+exp(\beta(e_{i1}-\phi_{out}))} \quad \text{(S3-10).}$$

An in-tether of the same size must behave the same, only the sign of the inclusions' energies on the tether flips, as it is now oppositely curved, and we have:

$$S_{in,i} = \frac{N_{i1}/M_1}{\rho_{i,global}} = \frac{M_1+M_2}{M_1}\frac{N_{i1}}{N_i} = \frac{M_1+M_2}{M_1}\frac{1}{1+exp(\beta(-e_{i1}-\phi_{in}))} \quad \text{(S3-11).}$$

Both $\phi_{out}$ and $\phi_{in}$ must be recovered by solving (S3-8) for each system, switching $e_{i1}$ to $-e_{i1}$ in the equation for $\phi_{in}$. For the ratio of sorting to an in-tether vs. to an out-tether relative to the global density for a species with curvature $c_{0,i}$ this approach yields (eq. 5, main text):

$$S_{in}/S_{out}(c_{0,i}) = (1 + exp(\beta(-2\kappa c_{0,i}\,\frac{a_0}{R} - \phi_{out})))/(1 + exp(\beta(2\kappa c_{0,i}\,\frac{a_0}{R} - \phi_{in}))) \quad \text{(S3-12)}$$

and is independent of the size of the tethered vs. flat regions. $R$ is the radius of the tether (Eq. 2 main text), assuming an idealized in- and out-tether.

To use this measure to extract e.g. $c_{0,i}$ of a specific protein species on the plasma membrane from experiments, one would ideally use knowledge of the underlying curvature distribution of all constituents of the membrane to solve eq. (S3-12) for $c_{0,i}$ after measuring $S_{in}/S_{out}(protein\ with\ c_{0,i})$. However, the curvature distribution is usually elusive.
Yet, in the following we will show that $\phi_{in/out}$ and hence the sorting ratio itself are approximately independent of the underlying inclusion type/ curvature distribution in many biologically relevant regimes. Therefore, measuring $S_{in}/S_{out}(protein\ with\ c_{0,i})$ indeed allows to solve eq. (S3-12), or for more realism, to compare simulated vs. experimental sorting, even when the experimental distribution is not known.

To show this, let us look at two arbitrary curvature distributions $\alpha, \beta$ on membrane tethers of the same size. To determine the four Lagrange multipliers $\phi^{\alpha}{}_{in/out}$ and $\phi^{\beta}{}_{in/out}$ we need to solve (S3-8) for each of them. Let us consider the out-tether (the in-tether can be solved in the same way). Then, for any two distributions with species of abundance $N_i{}^{\alpha}$, $N_i{}^{\beta}$:

$$M_1 = \sum_{i=0}^{I} \frac{N_i{}^{\alpha}}{1+exp(\beta(2\kappa\frac{c^{\alpha}{}_{0,i}}{R}a_0-\Phi^{\alpha}{}_{out}))} = \sum_{i=0}^{I}\frac{N_i{}^{\alpha}}{1+exp(\beta(D^{\alpha}{}_i-\Phi^{\alpha}{}_{out}))} \quad \text{(S3-13)}$$

$$M_1 = \sum_{i=0}^{I} \frac{N_i{}^{\beta}}{1+exp(\beta(2\kappa\frac{c^{\beta}{}_{0,i}}{R}a_0-\Phi^{\beta}{}_{out}))} = \sum_{i=0}^{I}\frac{N_i{}^{\beta}}{1+exp(\beta(D^{\beta}{}_i-\Phi^{\beta}{}_{out}))} \quad \text{(S3-14)}$$

The assumption we will need to make here, is that the curvatures induced on each membrane element $c_{0,i}\ a_0$ are relatively small in comparison to the tether radius $R$, which is realistic for our systems. We then can use a taylor expansion of each term in the above sums around small $D_i$, i.e. at $D_i = 0$. For simplicity let us set $\beta = k_BT = 1$. We expand to first (linear) order, giving:

$$\frac{N_i}{1+exp(D_i-\phi_{out})} = N_i(\frac{1}{1+exp(-\Phi_{out})} - D_i\frac{1}{1+exp(\Phi_{out})}(1-\frac{1}{1+exp(\Phi_{out})})) + O(D_i{}^2).$$

Setting (S3-13)=(S3-14) and using this approximation on both sides, it follows:

$$\frac{N^{\alpha}{}_{tot}}{1+exp(-\Phi^{\alpha}{}_{out})} - \sum_{i=0}^{I} N_i{}^{\alpha} D_i{}^{\alpha} \left(\frac{1}{1+exp(\Phi^{\alpha}{}_{out})} - \frac{1}{(1+exp(\Phi^{\alpha}{}_{out}))^2}\right)$$

$$\approx \frac{N^{\beta}{}_{tot}}{1+exp(-\Phi^{\beta}{}_{out})} - \sum_{i=0}^{I} N_i{}^{\beta} D_i{}^{\beta} \left(\frac{1}{1+exp(\Phi^{\beta}{}_{out})} - \frac{1}{(1+exp(\phi^{\beta}{}_{out}))^2}\right) \text{ (S3-15).}$$

From the zeroth order $\frac{N^{\alpha}{}_{tot}}{1+exp(-\Phi^{\alpha}{}_{out})} \approx \frac{N^{\beta}{}_{tot}}{1+exp(-\Phi^{\beta}{}_{out})}$we already know that the difference in $\phi_{out}$ between the two distributions $\Delta$ must be small, where $\Phi^{\alpha}{}_{out} = \Phi^{\beta}{}_{out} - \Delta$ .
To quantify the difference in more detail, we can hence linearize a second time, using a taylor expansion around small $\Delta$:

$$\frac{1}{1+exp(-\Phi^{\alpha}{}_{out})} \approx \frac{1}{1+exp(-\Phi^{\beta}{}_{out})} + \Delta\left(\frac{1}{1+exp(\Phi^{\beta}{}_{out})} - \frac{1}{(1+exp(\Phi^{\beta}{}_{out}))^2}\right) \text{ and}$$

$$\frac{1}{1+exp(\Phi^{\alpha}{}_{out})} - \frac{1}{(1+exp(\Phi^{\alpha}{}_{out}))^2} \approx \frac{1}{1+exp(\Phi^{\beta}{}_{out})} - \frac{1}{(1+exp(\Phi^{\beta}{}_{out}))^2} + \Delta O(D_i)$$

$$\approx \frac{1}{1+exp(\Phi^{\beta}{}_{out})} - \frac{1}{(1+exp(\Phi^{\beta}{}_{out}))^2},$$

where the linear term in $\Delta$ was dropped, since it will be second order in $\Delta, D_i$ once plugged into (S3-15) and we assume both are small.

Filling these second linearizations back into our first approximation (S3-15) we have:

$$\frac{N^{\beta}{}_{tot}}{1+exp(-\Phi^{\beta}{}_{out})} - \sum_{i=0}^{I} N_i{}^{\beta} D_i{}^{\beta} \left(\frac{1}{1+exp(\Phi^{\beta}{}_{out})} - \frac{1}{(1+exp(\phi^{\beta}{}_{out}))^2}\right)$$

$$\approx \frac{N^{\alpha}{}_{tot}}{1+exp(-\Phi^{\beta}{}_{out})} + \Delta\left(\frac{N^{\alpha}{}_{tot}}{1+exp(\Phi^{\beta}{}_{out})} - \frac{N^{\alpha}{}_{tot}}{(1+exp(\Phi^{\beta}{}_{out}))^2}\right)$$

$$- \sum_{i=0}^{I} N_i{}^{\alpha} D_i{}^{\alpha} \left(\frac{1}{1+exp(\Phi^{\beta}{}_{out})} - \frac{1}{(1+exp(\Phi^{\beta}{}_{out}))^2}\right)$$

Using the fact that both distributions each fully cover their tether and hence $N^{\beta}{}_{tot} = N^{\alpha}{}_{tot} = N_{tot}$ this can be rearranged to:

$$\Delta\left(\frac{N_{tot}}{1+exp(\Phi^{\beta}{}_{out})} - \frac{N_{tot}}{(1+exp(\Phi^{\beta}{}_{out}))^2}\right) \approx \left(\frac{1}{1+exp(\Phi^{\beta}{}_{out})} - \frac{1}{(1+exp(\Phi^{\beta}{}_{out}))^2}\right) \sum_{i=0}^{I}(N_i{}^{\alpha} D_i{}^{\alpha} - N_i{}^{\beta} D_i{}^{\beta})$$

(S3-16).

Hence:

$$\Phi^{\beta}{}_{out} - \Phi^{\alpha}{}_{out} = \Delta \approx \frac{1}{N_{tot}} \sum_{i=0}^{I}(N_i{}^{\alpha} D_i{}^{\alpha} - N_i{}^{\beta} D_i{}^{\beta}) = mean(D^{\alpha}{}_i) - mean(D^{\beta}{}_i) \text{ (S3-17).}$$

Therefore, for any two curvature distributions covering the membrane, the sorting ratio of an individual species, $S_{in}/S_{out}(c_{0,i})$, in both distributions will be the same to linear order, as long as the two distributions have the same mean curvature. This approximation appears to be very precise for curvatures considered in this work with $0.3 < |c_{0,i}/R|$ as there are hardly detectable differences in sorting ratios even for strongly asymmetric distributions as visualized in Supplementary Figure S14b.

As we show in the main text, the mean of a curvature distribution on the plasma membrane can e.g. be measured via the force difference in combined in-out-tether pulling. Therefore, measuring the sorting ratio of specific protein on in- vs. out-tethers afterwards, and comparing it to simulated sorting of distributions with the same mean $< c_0 >$ allows to extract the curvature $c_{0,i}$ of that specific protein species.
